# ‘Q-niche’: topology-preserving niche hypervolumes built using a quantum mechanics modelling analogue

**DOI:** 10.64898/2026.10.29.762789

**Authors:** Joseph Thomas, Juliano Morimoto

## Abstract

Hutchinson’s *n*-dimensional niche hypervolume underlies many ecological analyses, yet estimating hypervolumes from finite occurrence data remains challenging, particularly when niches contain non-convex structure such as separated regions or internal holes. Here, we introduce Q-niche, a quantum-mechanics-inspired algorithm for ecological hypervolume estimation. Using an inverse Schrödinger formulation, Q-niche transforms an occurrence-derived Gaussian field into a potential surface whose low-potential regions define a hypervolume represented by a stochastic point cloud. We evaluated Q-niche against Gaussian kernel density estimation and one-class support vector machine hypervolume estimators using synthetic niches, virtual species and empirical *Acacia* occurrence data. Across controlled synthetic benchmarks, Q-niche preserved imposed topological features more consistently than either alternative and showed closer overall topological agreement with occurrence clouds. This preservation remained robust across the tested parameter settings. Virtual-species analyses showed that stronger preservation of environmental topology did not necessarily produce greater overlap with the known geographical distribution, showing that topology and geographical range are not necessarily coupled. The empirical *Acacia* dataset revealed that the inferred size and topology of a niche hypervolume is estimator-dependent, favouring the use of Q-niche when hypervolume topology is the focus of research due to its topology preservation performance. Overall, Q-niche provides a practical tool for reconstructing complex niche hypervolumes and investigating how their geometry and topology relate to potential geographical distributions.

## Introduction

The ecological niche is one of the most used and debated concepts in ecology [1, 2, 3]. A particularly useful formulation of the niche in both theoretical and applied ecology is that of the Hutchinsonian niche, which is represented as an *n*-dimensional hypervolume whose axes describe environmental conditions relevant to species persistence [2, 4, 5, 6]. The Hutchinsonian niche hypervolume concept has become central to quantitative ecology, informing theories of resource partitioning and species coexistence [7, 8, 9, 10], as well as approaches linking species’ environmental requirements to their geographical distributions in biogeography and macroecology [2, 11, 12]. A major reason for the widespread use of the niche hypervolume concept is its versatility. Approaches relying on hypervolumes provide a general framework for quantifying niche breadth, position and overlap, phenotypic and functional trait spaces and the functional diversity of biological assemblies [13, 14, 15, 16, 17]. Therefore, the Hutchinsonian niche hypervolume concept is fundamental to ecology.

Recent studies have associated the geometric and topological features of the niche hypervolume with hypotheses of species’ responses to their environment and to each other. Consequently, there has been increasing interest in understanding how the geometry and topology of niche hypervolumes inform ecological and evolutionary properties of species. For instance, Blonder proposed that holes within ecological hypervolumes may themselves contain ecological information [12], for example by reflecting exclusion or susceptibility to invasion. More broadly, ecological theory has long recognised that realised niches need not form simple convex regions. Maguire described the realised niche as a potentially smaller and differently shaped subset of environmental space [18], while subsequent work showed that realised responses along environmental gradients may be skewed, non-Gaussian or bimodal [19, 20]. Recent work has begun to quantify such geometric and topological properties directly and to investigate their ecological and evolutionary significance. For example, Caron et al. quantified both niche hypervolume size and holes across 418 gymnosperm species and found contrasting evolutionary patterns: niche volume showed moderate phylogenetic signal, whereas hole structure showed little phylogenetic constraint [21].

The theoretical suppositions and empirical null results can signify that the topology of niche hypervolumes is unimportant, or that our ability to generate topologically-preserving hypervolumes needs advancement. Given that topology has only recently become the focus of niche theory [22], the latter seems likely. Established hypervolume estimators address the latter in different ways. Range boxes, ellipsoidal representations and convex hulls provide relatively simple geometrical descriptions of an occurrence cloud, but constrain the shapes that can be reconstructed from the observations [14, 23, 24, 25]. More flexible approaches include Gaussian kernel density estimation (KDE) and one-class support vector machines (SVM), both of which can delineate non-convex or disconnected regions in multidimensional space [16, 17]. These methods nevertheless encode different rules for assigning unsampled environmental conditions to the estimated niche: KDE delineates level sets of an occurrence-derived density surface, whereas a one-class SVM estimates a binary support boundary around the observations [17]. Consequently, estimators fitted to the same occurrence data will differ not only in overall niche extent and overlap, but also in whether separated components or internal gaps are retained or interpolated across [12, 17, 26]. Estimating a niche hypervolume from occurrence data is therefore an inferential problem: a finite sample of observed environmental conditions must be used to determine the extent and internal structure of a continuous region that also contains unsampled environmental space [16, 17]. Measures such as hypervolume size and Jaccard overlap quantify important aspects of overall geometrical agreement, but do not explicitly describe connected components or internal topological structure [12, 22, 26]. Therefore, our knowledge of how established estimators deal with, distort, or preserve the geometry and topology of niche hypervolumes is limited. Some estimators impose convexity on the modelled niche [14, 23, 24, 25], yet realised niches may be non-convex, may contain separated regions or include internal gaps [26]. Many related and unrelated processes may give rise to these geometric and topological features, such as environmental availability, historical constraints or patchy or biased sampling [2, 12, 18, 19, 20, 21, 27]. While no estimator can resolve all of these empirical limitations, understanding how different algorithms preserve geometric and topological structure is necessary before such properties can be interpreted ecologically or evolutionarily. If our estimators lack the ability to preserve topology and geometry of hypervolumes well, then we lack the fundamental knowledge and tools for testing existing ecological hypotheses related to niche hypervolume topology.

Here, we introduce ‘Q-niche’ – a quantum mechanics-inspired algorithm to estimate niche hypervolumes. This approach draws on quantum-mechanics-inspired methods for data analysis, specifically on the quantum clustering framework developed by Horn and Gottlieb [28, 29]. The use of quantum-inspired modelling outside physics is not unprecedented in ecology: related ideas have been explored in population dynamics and localisation [30, 31], persistence under environmental change [32], and uncertainty in species distributions [33]. Broader analogies between ecological and quantum principles have also been proposed [34], alongside more recent quantum-computing approaches to ecological problems [35]. In the Q-niche approach developed here, quantum clustering treats a Gaussian kernel field constructed from observation data points as the ground-state wavefunction in a time-independent Schrödinger equation. Rather than specifying a potential and solving for the wavefunction, as in the conventional quantum mechanics problem, the relationship is inverted so that the observed data define the wavefunction and the corresponding potential landscape is derived [28, 29, 36]. Low-potential regions tend to occur around concentrations of observation points, whereas higher-potential regions can occur farther from observations. Classic quantum clustering uses this potential landscape to characterise structure within observed data. Q-niche adapts this approach and uses a potential sublevel set to define the support of a continuous ecological hypervolume, together with a stochastic representation that permits estimation of volume, overlap, topology and geographical projection (Fig. 1).

**Figure 1.**
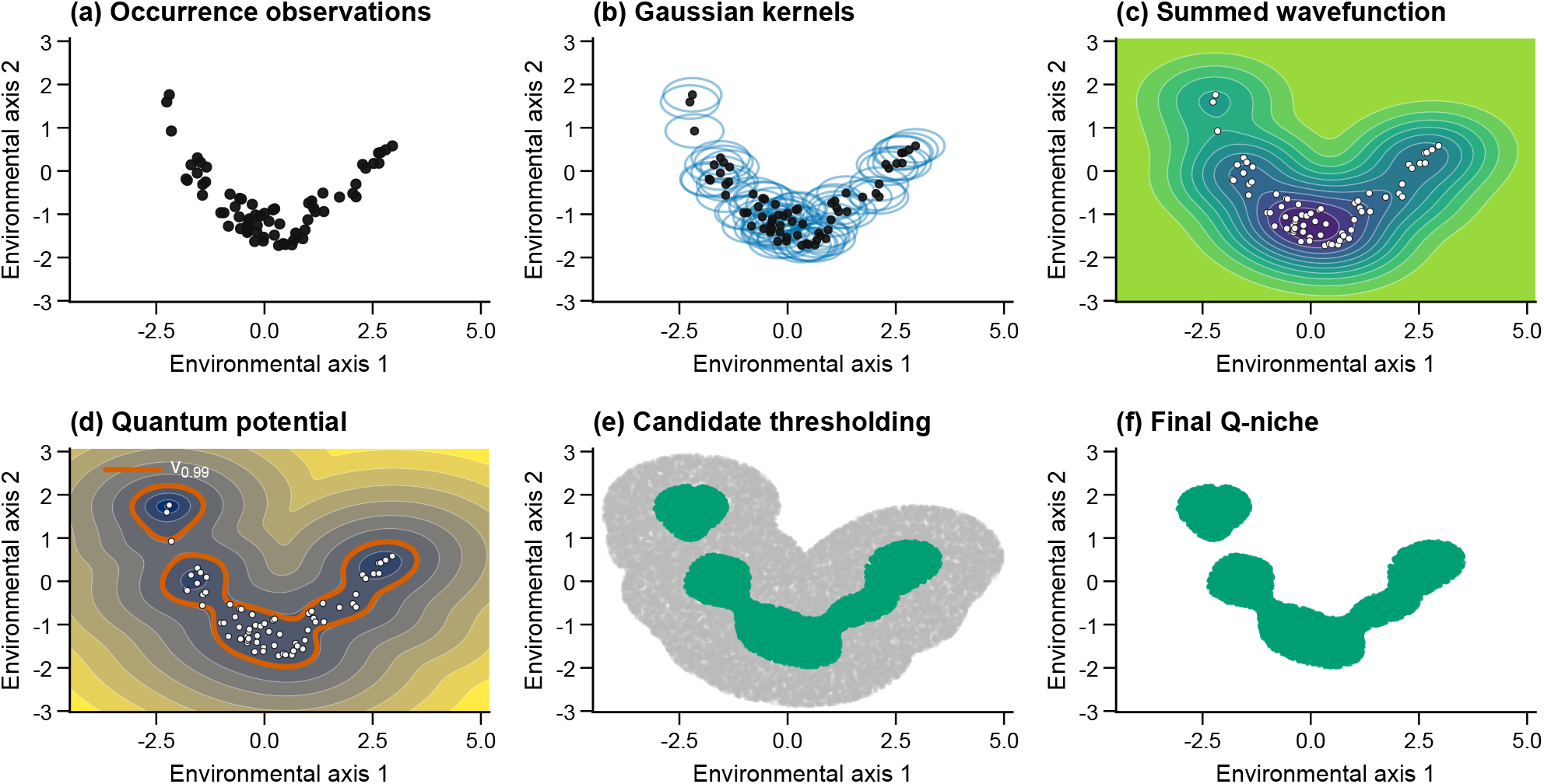
Conceptual workflow for constructing a Q-niche hypervolume. (a) Occurrence observations in a two-dimensional environmental space. (b) A Gaussian function is centred on each occurrence, with spread determined by the isotropic bandwidth *h*. (c) The Gaussian functions are summed to form the occurrence-derived field *ψ*_*h*_(**x**). (d) The inverse Schrödinger formulation transforms this field into the quantum potential *V*_*h*_(**x**). Contours indicate equal potential values; the orange contour denotes *V*_*h*_(**x**) = *v*_0.99_, where *v*_0.99_ is the 99th percentile of potential values evaluated at the occurrence observations. For a fixed potential surface and candidate region, lowering the threshold reduces the retained niche region, whereas raising it expands the region. (e) Candidate points are sampled from the finite region surrounding the observations. Points satisfying *V*_*h*_(**z**_*j*_) *≤ v*_0.99_ are retained and rejected points are shown in grey. (f) The retained points form the stochastic point-cloud representation of the final Q-niche hypervolume.

To evaluate the broader utility of Q-niche, we first describe the conceptual and mathematical basis of the method and then evaluate its performance against Gaussian KDE and one-class SVM hypervolume estimators [16, 17]. We then use two- and three-dimensional synthetic niches to test the ability of Q-niche, Gaussian KDE, and SVM to recover known geometry and topology. Third, we simulate virtual species with known environmental and geographical distributions to test how Q-niche performs relative to the established algorithms in an ecological setting for which the true geographical distribution is known. Finally, we apply and compare the estimators using occurrence data from five Australian *Acacia* species for which the true niche and distribution are unknown. Across these analyses, we compare hypervolume size, position and overlap, use persistent homology to quantify preservation of topological structure, and examine how differences among environmental hypervolume estimates affect potential geographical projections. Together, these analyses establish Q-niche as a new approach to ecological hypervolume estimation and test its ability to preserve complex niche geometry and topology.

## Q-niche: the model

### Conceptual formulation

Quantum potential niche hypervolumes (‘Q-niche’) estimate an occurrence-based ecological niche hypervolume from observations represented in continuous environmental space. Each occurrence is first associated with a Gaussian function whose spread is controlled by a single isotropic bandwidth applied equally across environmental dimensions, and these functions are summed to form a smooth occurrence-derived Gaussian field, as in Gaussian kernel density estimation [16, 17]. Q-niche then applies the inverse Schrödinger formulation used in quantum clustering: the Gaussian field takes the role of the wavefunction, from which an associated quantum potential is derived [28, 29].

The resulting potential provides a transformed representation of the occurrence data. Low-potential regions are generally associated with environmental conditions locally supported by the observations, whereas higher-potential regions can occur farther from the observations or between separated regions of support. Q-niche defines the estimated niche by retaining environmental conditions below an occurrence-derived potential threshold. The potential quantile is specified by the investigator; throughout the main analyses we use *q* = 0.99, so that the resulting threshold is the 99th percentile of the occurrence-point potentials. Points are then sampled uniformly from a finite region surrounding the observations and retained when their potential falls below this threshold. The retained points provide a stochastic representation of the estimated niche region: unlike the original occurrence points, which are the observations used to fit the model, these newly sampled points represent environmental conditions classified by Q-niche as belonging to the niche. The complete construction is illustrated in Fig. 1. The following sections describe this construction and the resulting hypervolume estimator in detail; a full derivation from the Schrödinger equation, further quantum-mechanical background, and details of the bandwidth calculation are provided in Appendix A. Note that Q-niche is *quantum-inspired* rather than a physical model of ecological systems. The inverse Schrödinger formulation is used as a mathematical transformation of the occurrence-derived Gaussian field. The resulting potential is not interpreted as physical energy, occurrence probability or physiological suitability. This differs from the classical use of these concepts in physics, but as shown here, provides a useful framework for modelling niche hypervolumes.

### From occurrence data to a quantum potential

Let

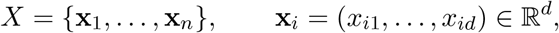

denote a set of *n* occurrence observations represented in a *d*-dimensional environmental space, where **x**_*i*_ is the environmental coordinate vector associated with occurrence *i*.

Q-niche uses a single isotropic Gaussian bandwidth *h >* 0, such that the bandwidth vector is

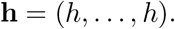

The bandwidth is estimated from local nearest-neighbour distances using a procedure adapted from Nasios and Bors [36]. For a dataset containing *n* occurrence observations, we define *K* as the number of neighbouring occurrences used to characterise the local spacing around each observation, and set

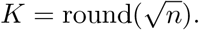

For each occurrence **x**_*i*_, we identify its *K* nearest other occurrences, denoted **x**_*i*(1)_, … , **x**_*i*(*K*)_, and calculate the local squared-distance statistic

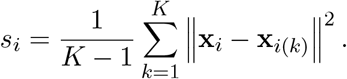

These local values are then averaged across all *n* occurrences, and the bandwidth is defined as the square root of this average,

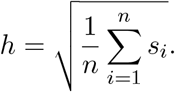

The resulting value *h* is applied equally across all environmental dimensions. Further details of the bandwidth construction and its relationship to the approach of Nasios and Bors are provided in Appendix A.

For an arbitrary environmental coordinate **x** = (*x*_1_, … , *x*_*d*_), *x*_*j*_ denotes its coordinate in dimension *j*, while *x*_*ij*_ denotes the coordinate of occurrence *i* in the same dimension. We denote the squared distance between **x** and occurrence *i* in bandwidth-standardised coordinates by 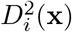, such that

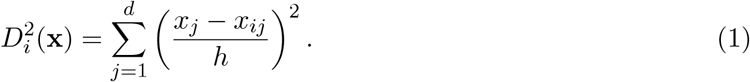

Each occurrence contributes a Gaussian function centred on its environmental coordinates (Fig. 1b). Summing these functions (Fig. 1c) gives the occurrence-derived Gaussian field

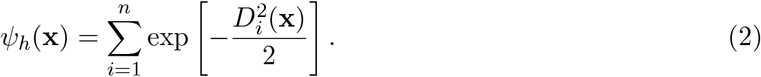

Apart from a multiplicative normalising constant, this field has the form of a Gaussian kernel sum [17].

Following quantum clustering, *ψ*_*h*_ is treated mathematically as the ground-state wavefunction of a time-independent Schrödinger equation [28, 29]. In the conventional physics formulation, the potential is specified, and the wavefunction is solved for; here the occurrence-derived field is known, and the relationship is instead rearranged to obtain its associated potential:

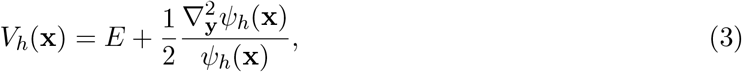

where **y** denotes the bandwidth-standardised environmental coordinates, 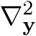 is the corresponding Laplacian (the sum of second derivatives across dimensions), and *E* is an additive constant. For the Gaussian field in Equation 2, this expression can be evaluated analytically to give

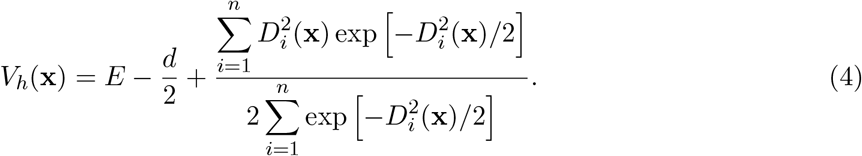

The derivation from the time-independent Schrödinger equation, including the role of the bandwidth-standardised coordinates, is given in Appendix A. Equation 4 provides the quantity *V*_*h*_(**x**) used directly by Q-niche (Fig. 1d). At any environmental coordinate, the potential is determined by one half of a Gaussian-weighted mean of its squared bandwidth-standardised distances from the occurrence observations, together with the constant shift *E −d/*2. The resulting surface forms a continuous landscape in which locally supported regions tend to have lower potential and separated regions can be divided by higher-potential barriers. Unlike a conventional KDE, Q-niche therefore delineates the hypervolume from the transformed potential rather than directly from the magnitude of the Gaussian density field. The constant *E* shifts all potential values equally and therefore does not affect their ordering or the sublevel sets used to define Q-niche; we set *E* = 0 throughout.

### Hypervolume delineation and stochastic representation

The quantum potential is defined throughout environmental space, but evaluating an unbounded space is neither necessary nor computationally feasible. Q-niche therefore first defines a finite candidate region around the occurrence observations. Following the stochastic sampling framework of the `hypervolume R` package [17], this region is the union of occurrence-centred hyperspheres,

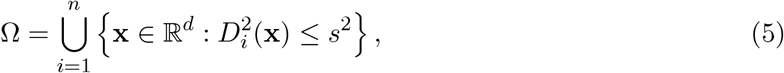

where *s* controls the extent of candidate sampling relative to the Gaussian bandwidth. We used *s* = 3, corresponding to `sd.count` = 3 in the `hypervolume` package, so that each hypersphere had radius 3*h*. This parameter sets the maximum region available to the estimator; within this region, the final Q-niche boundary is determined by the potential threshold described below. Candidate points were sampled uniformly from Ω using the sampling procedure implemented by `hypervolume` [17]. This procedure accounts for overlap among neighbouring sampling regions when constructing the candidate point cloud. The baseline stochastic representation used `samples.per.point` = 100. The quantum potential *V*_*h*_ was evaluated at both the occurrence observations and the candidate points. The threshold introduced in the conceptual formulation and illustrated in Fig. 1d is calculated from the occurrence-point potentials. For a chosen quantile *q*, its value is

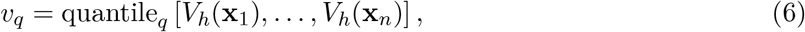

where quantile_*q*_ denotes the empirical *q*th quantile. We used *q* = 0.99 in the main analyses. Candidate points within Ω were then retained when their potential was at or below *v*_*q*_ (Fig. 1e), producing the final hypervolume point cloud (Fig. 1f). Thus, *s* controls the region from which candidate points are sampled, whereas *v*_*q*_ determines which of those points are retained. The estimated Q-niche hypervolume is the low-potential region

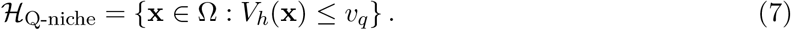

In practice, this region was represented by Monte Carlo sampling within Ω. The potential was evaluated at each uniformly sampled candidate point, and points with *V*_*h*_(**x**) *≤ v*_*q*_ were retained, whereas points above the threshold were discarded. The retained candidate points therefore provide a stochastic representation of the Q-niche sublevel set. Because the candidate points are sampled uniformly from Ω, hypervolume size is estimated from the fraction satisfying the Q-niche criterion. If *M* candidate points are generated and *M*_Q-niche_ are retained,

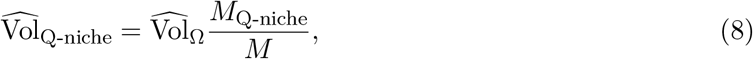

where 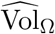 is the estimated volume of the candidate region. The occurrence observations and retained Q-niche points were stored separately within each `Hypervolume` object: the former remained the input data, whereas the latter formed the stochastic representation of the estimated niche used for downstream analyses. The associated volume and point density were also stored in the object, allowing Q-niche estimates to be analysed using existing visualisation, inclusion and set-operation functions in the `hypervolume` workflow.

## Results

### Q-niche is more consistent at preserving known topology across synthetic datasets

We first evaluated Q-niche against Gaussian KDE and one-class SVM hypervolumes [17] using synthetic occurrence clouds whose underlying topology was known exactly. Five two-dimensional regions were considered: a spiral band, filled ellipse, annulus, concave banana and two separated disks. Three-dimensional tests comprised a solid ball, solid torus and hollow spherical shell. Point clouds, representing occurrence points, were sampled from each region at nested sample sizes of *n* = 300, 900 and 1500, and the resulting point clouds were used to construct each hypervolume estimate. Representative two-dimensional reconstructions are shown in Fig. 2. Generating definitions, parameter values, analytical areas or volumes and expected topological structure for all eight niches are given in Table S1.

**Figure 2.**
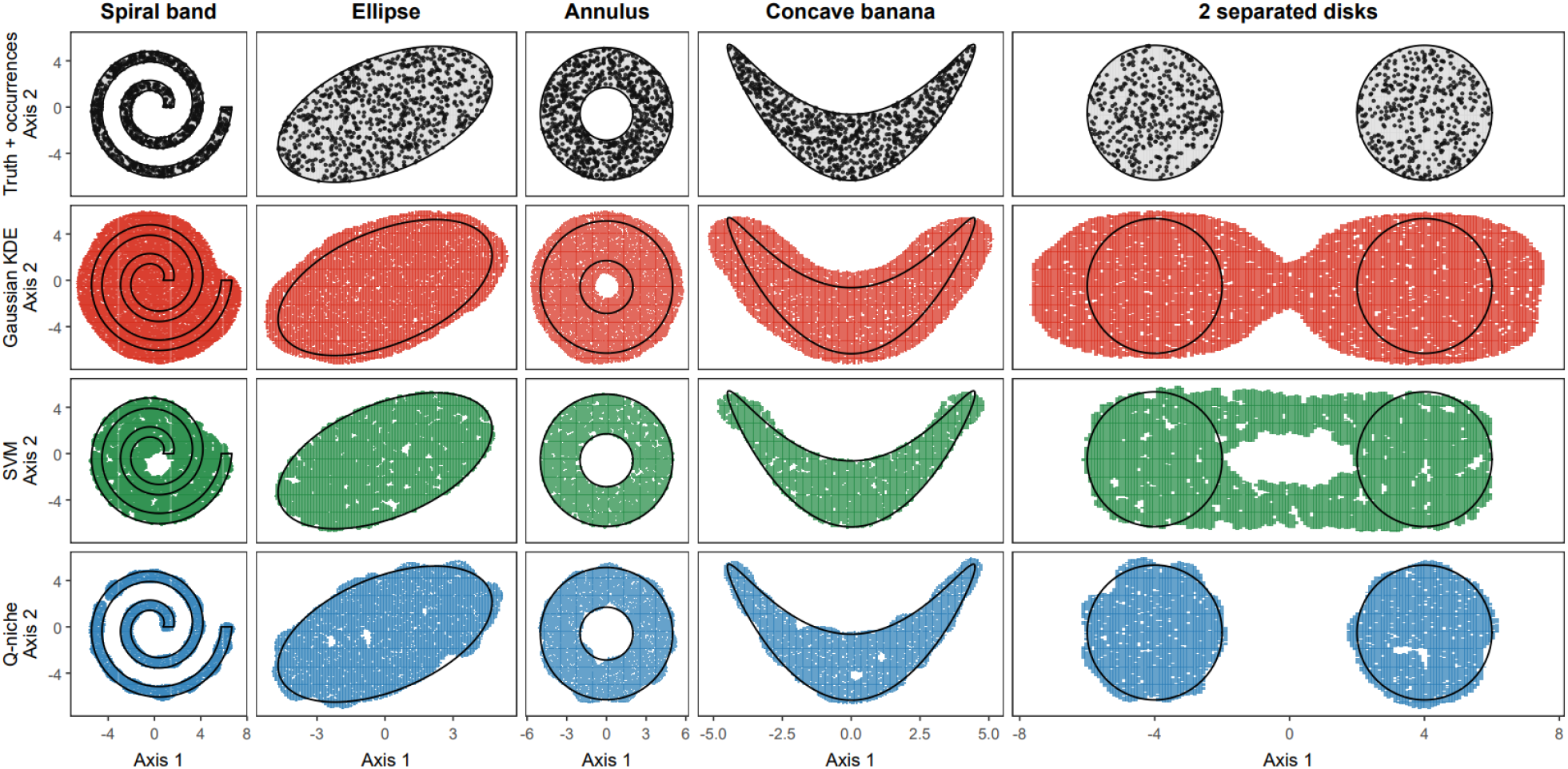
Representative reconstruction of known two-dimensional synthetic niches by alternative hypervolume estimators. Five analytically defined synthetic niches were reconstructed using Gaussian kernel density estimation (KDE), one-class support vector machines (SVM) and Q-niche. A representative occurrence sample size of *n* = 900 is shown. The upper row shows the occurrence observations and analytical niche boundaries; subsequent rows show the stochastic hypervolume representations produced by each estimator under the baseline parameterisation. Black lines indicate the known niche boundaries.

Persistent homology was calculated for the occurrence and reconstructed hypervolume point clouds (representative two-dimensional examples are shown in Fig. 2), with *H*_0_, *H*_1_ and, for the three-dimensional niches, *H*_2_ describing connected components, loops and enclosed voids, respectively (see Methods). This approach measures how long topological features remain detectable across distance scales, allowing their persistence to be compared between point clouds [37, 38]. For computational tractability, we analysed 10 reproducible samples of 300 points from each source cloud and recorded the maximum persistence in each homology dimension. To assess whether these features were more persistent than those arising in a simple convex region, we generated 20 reference point clouds from filled ellipses or ellipsoids. Each reference contained 300 points and matched the centroid and covariance of the full source cloud. For each homology dimension, we asked whether the maximum persistence in each observed sample exceeded the largest maximum persistence across the 20 reference point clouds. We report the fraction of the ten samples meeting this criterion as the exceedance proportion: a value of 1 means all ten exceeded the reference criterion, whereas 0 means none did. For the deliberately non-trivial niches, high exceedance proportions in both the occurrence and reconstructed clouds indicate consistent expression of the expected topological feature, whereas low proportions are expected for the nominally topologically simple controls. Topological similarity was also quantified using bottleneck distance, normalised by the diameter of the corresponding occurrence cloud, so that smaller values indicate closer agreement with the observed topological structure.

Across the five two-dimensional niches and all three sample sizes, Q-niche had the smallest overall mean normalised bottleneck distance in both *H*_0_ and *H*_1_ (Table S2). The deliberately non-trivial niches showed the clearest differences among estimators (Table 1). The annular *H*_1_ feature exceeded the matched ellipsoid criterion in every Q-niche and SVM repetition at all three sample sizes, whereas Gaussian KDE exceedance increased with sample size. For the two separated disks, Q-niche reproduced the expected finite *H*_0_ structure in every repetition at every sample size. In contrast, SVM showed no exceedance, while Gaussian KDE exceeded the criterion only occasionally at the largest sample size.

**Table 1:**
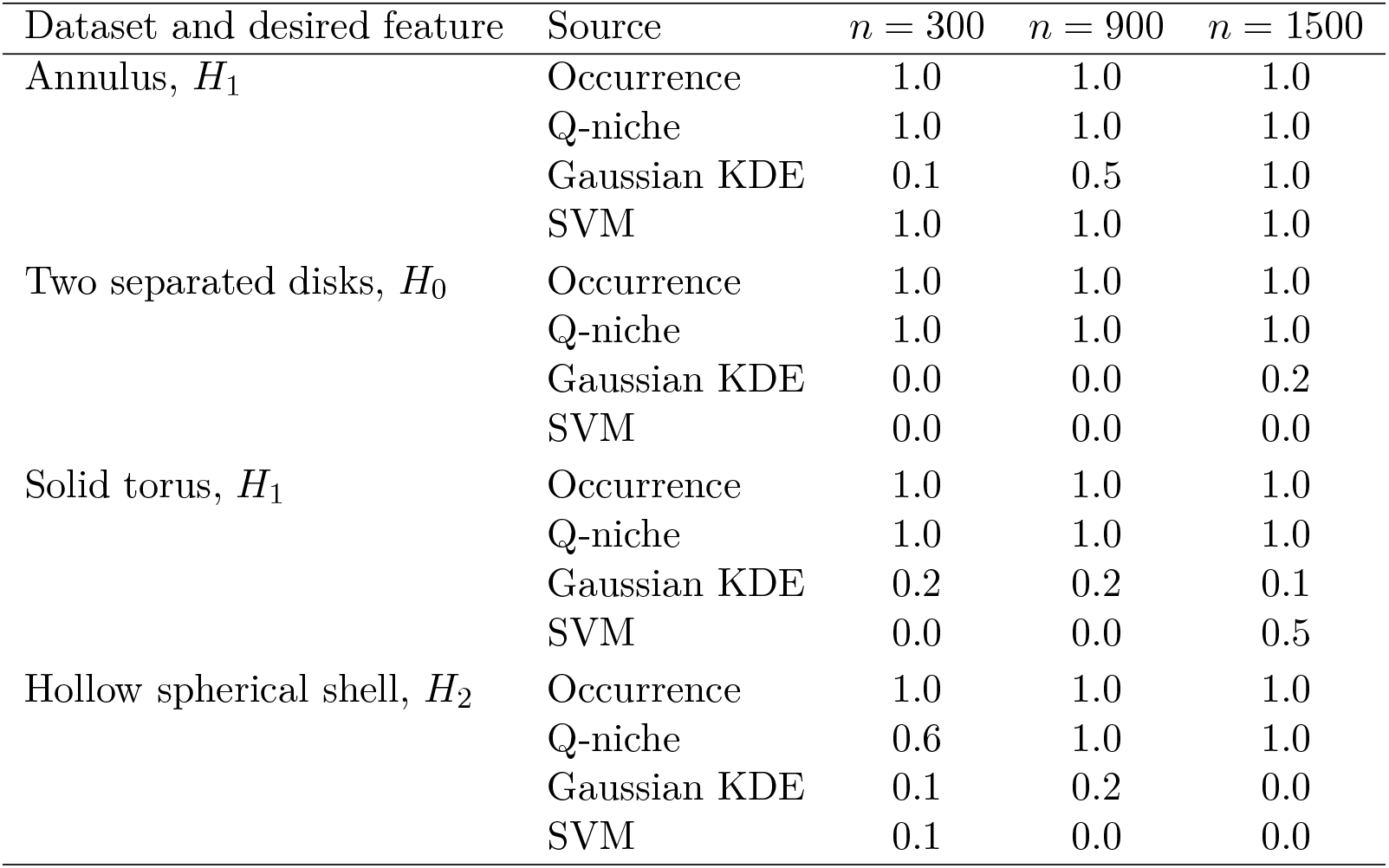
Exceedance proportions for the expected non-trivial topological features of the synthetic niches. Values show how consistently each expected feature exceeded a reference criterion based on finite samples from filled ellipses or ellipsoids. For each source and occurrence sample size, 20 reference clouds of 300 points were matched to the full source cloud’s centroid and covariance. The largest finite persistence across these references defined the criterion for each homology dimension. Values give the proportion of ten 300-point repetitions whose maximum persistence exceeded that criterion: 1 indicates exceedance in every repetition and 0 indicates none. High values in both occurrence and reconstructed clouds support retention of the expected feature. *H*_0_, *H*_1_ and *H*_2_ describe connected components, loops and enclosed voids, respectively.

| Dataset and desired feature | Source | $n = 300$ | $n = 900$ | $n = 1500$ |
| --- | --- | --- | --- | --- |
| Annulus, $H_1$ | Occurrence | 1.0 | 1.0 | 1.0 |
|  | Q-niche | 1.0 | 1.0 | 1.0 |
|  | Gaussian KDE | 0.1 | 0.5 | 1.0 |
|  | SVM | 1.0 | 1.0 | 1.0 |
| Two separated disks, $H_0$ | Occurrence | 1.0 | 1.0 | 1.0 |
|  | Q-niche | 1.0 | 1.0 | 1.0 |
|  | Gaussian KDE | 0.0 | 0.0 | 0.2 |
|  | SVM | 0.0 | 0.0 | 0.0 |
| Solid torus, $H_1$ | Occurrence | 1.0 | 1.0 | 1.0 |
|  | Q-niche | 1.0 | 1.0 | 1.0 |
|  | Gaussian KDE | 0.2 | 0.2 | 0.1 |
|  | SVM | 0.0 | 0.0 | 0.5 |
| Hollow spherical shell, $H_2$ | Occurrence | 1.0 | 1.0 | 1.0 |
|  | Q-niche | 0.6 | 1.0 | 1.0 |
|  | Gaussian KDE | 0.1 | 0.2 | 0.0 |
|  | SVM | 0.1 | 0.0 | 0.0 |

The three-dimensional benchmarks extended the same comparison to loop and enclosed-void structure. At *n* = 900, Q-niche had lower mean normalised bottleneck distances than Gaussian KDE and SVM for the solid-torus *H*_1_ and hollow-shell *H*_2_ features (Table S2). Q-niche exceeded the matched ellipsoid criterion for the toroidal feature in every repetition at all three sample sizes. For the hollow shell, exceedance increased with sample size, reaching every repetition at *n* = 900 and 1500. Gaussian KDE and SVM showed substantially less consistent exceedance for both features (Table 1).

The nominally topologically simple controls showed comparatively little evidence of introduced persistent structure. Neither the occurrence clouds nor any estimator exceeded the *H*_1_ matched ellipsoid criterion for the concave banana at any sample size. Exceedance was rare for the filled ellipse, while Q-niche exceedance for the solid ball was rare in *H*_1_ and absent in *H*_2_ under the baseline parameterisation. The spiral band was less suitable as a simple negative control because its finite occurrence clouds frequently contained persistent *H*_1_ structure relative to the matched ellipsoid criterion, despite the generating region being simply connected. Complete results are provided in Table S2.

Across the sensitivity parameterisations at *n* = 900 (Fig. 3), Q-niche retained an exceedance proportion of 1.00 for the annulus *H*_1_, separated-disk *H*_0_, solid-torus *H*_1_ and hollow-shell *H*_2_ features, indicating robust preservation under the parameter changes examined. The corresponding ranges for Gaussian KDE and SVM were substantially wider for several of these features. Importantly, this robustness was accompanied by low exceedance in the topologically simple controls, indicating little evidence of introduced persistent structure.

**Figure 3.**
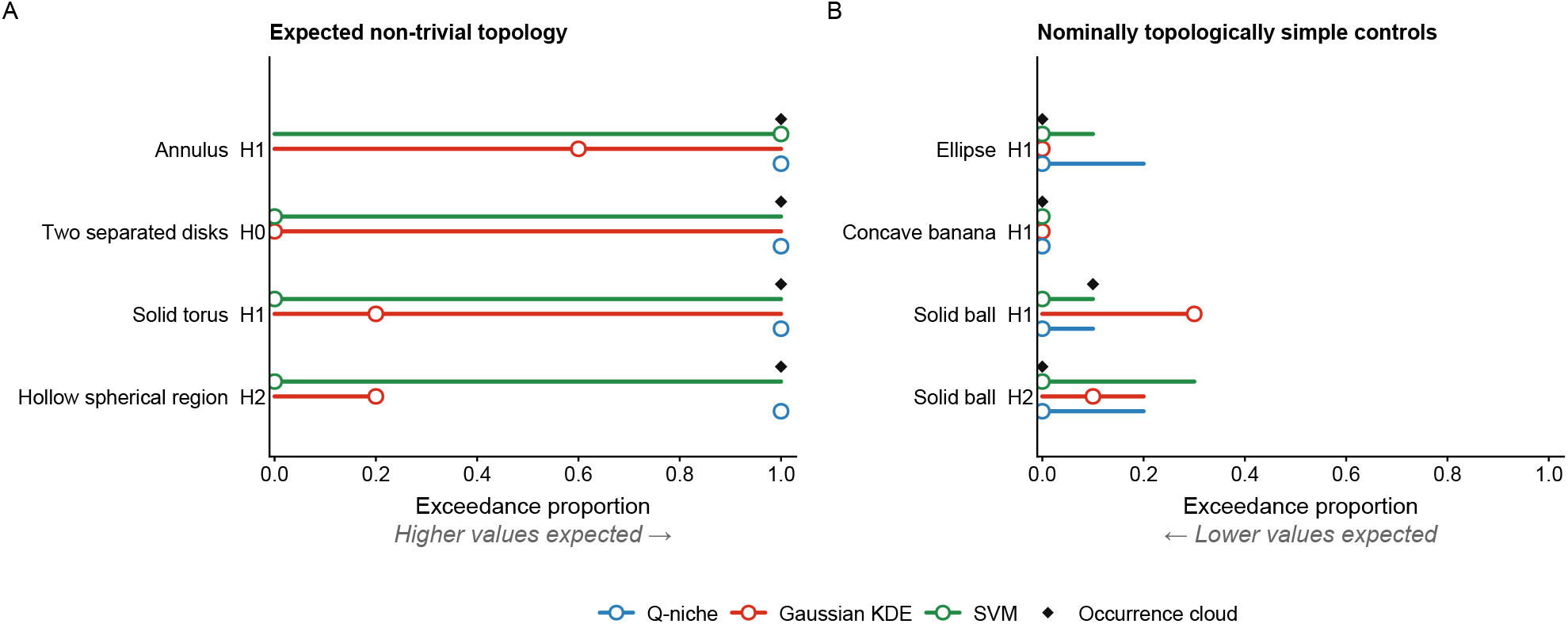
Sensitivity of synthetic niche topology to estimator parameterisation. Exceedance proportions relative to the matched ellipsoid criterion are shown at *n* = 900 for (A) the expected non-trivial topological features and (B) nominally topologically simple controls. Solid coloured lines show the minimum-to-maximum exceedance proportion across the tested sensitivity parameterisations for each estimator, and open circles indicate the corresponding baseline parameterisation. Filled black diamonds show the exceedance proportion of the occurrence cloud. Higher exceedance proportions are expected when the known features are retained in panel A, whereas lower proportions are expected for the controls without those features in panel B. Expected non-trivial features comprise the annulus (*H*_1_), two separated disks (*H*_0_), solid torus (*H*_1_) and hollow spherical shell (*H*_2_). Simple controls comprise the filled ellipse (*H*_1_), concave banana (*H*_1_), and solid ball (*H*_1_ and *H*_2_).

Repeating the two-dimensional analysis at *n* = 900 across independently generated occurrence clouds reproduced the same broad pattern: Q-niche strongly preserved the disconnected *H*_0_ structure of the separated disks, while both Q-niche and SVM showed low *H*_1_ bottleneck distances for the annulus (Table S3). Together, these benchmarks demonstrate Q-niche’s clear advantage in preserving disconnected components, loops and enclosed voids more consistently than Gaussian KDE or SVM. This preservation remained robust across the tested parameter settings, with little evidence of introduced persistent structure in the simple controls.

### Alternative estimators differ in recovery of known synthetic niche geometry

We next assessed geometrical recovery of the same eight synthetic niches used in the topological comparison, focusing on Jaccard similarity and relative area or volume error. Jaccard similarity measures the overlap between the estimated and known niche regions relative to their union, with higher values indicating closer agreement. Relative area or volume error expresses the difference between estimated and analytical niche size as a percentage of the analytical size; positive values indicate overestimation, negative values indicate underestimation, and values closer to zero indicate more accurate size estimates.

The three estimators produced markedly different geometrical reconstructions of the synthetic niches (Fig. 2; Table 2). In two dimensions, Q-niche performed particularly well for the spiral band and two separated disks. At *n* = 900, it showed substantially less area overestimation and greater Jaccard similarity with both known regions than SVM or Gaussian KDE. SVM more closely reconstructed the filled ellipse, annulus and concave banana, with smaller absolute relative area errors and higher Jaccard similarities than Q-niche at *n* = 900. Gaussian KDE consistently overestimated area across all five two-dimensional regions and all three occurrence sample sizes. The three-dimensional results showed the same dependence on niche geometry. At *n* = 900, Q-niche reconstructed the solid torus with less volume overestimation and higher Jaccard similarity than SVM or Gaussian KDE, whereas SVM most closely reproduced the solid ball and hollow spherical shell (Table 2). Thus, the estimator showing the closest geometrical agreement depended on niche structure and did not necessarily coincide with the estimator that most consistently preserved the expected topology.

**Table 2:**
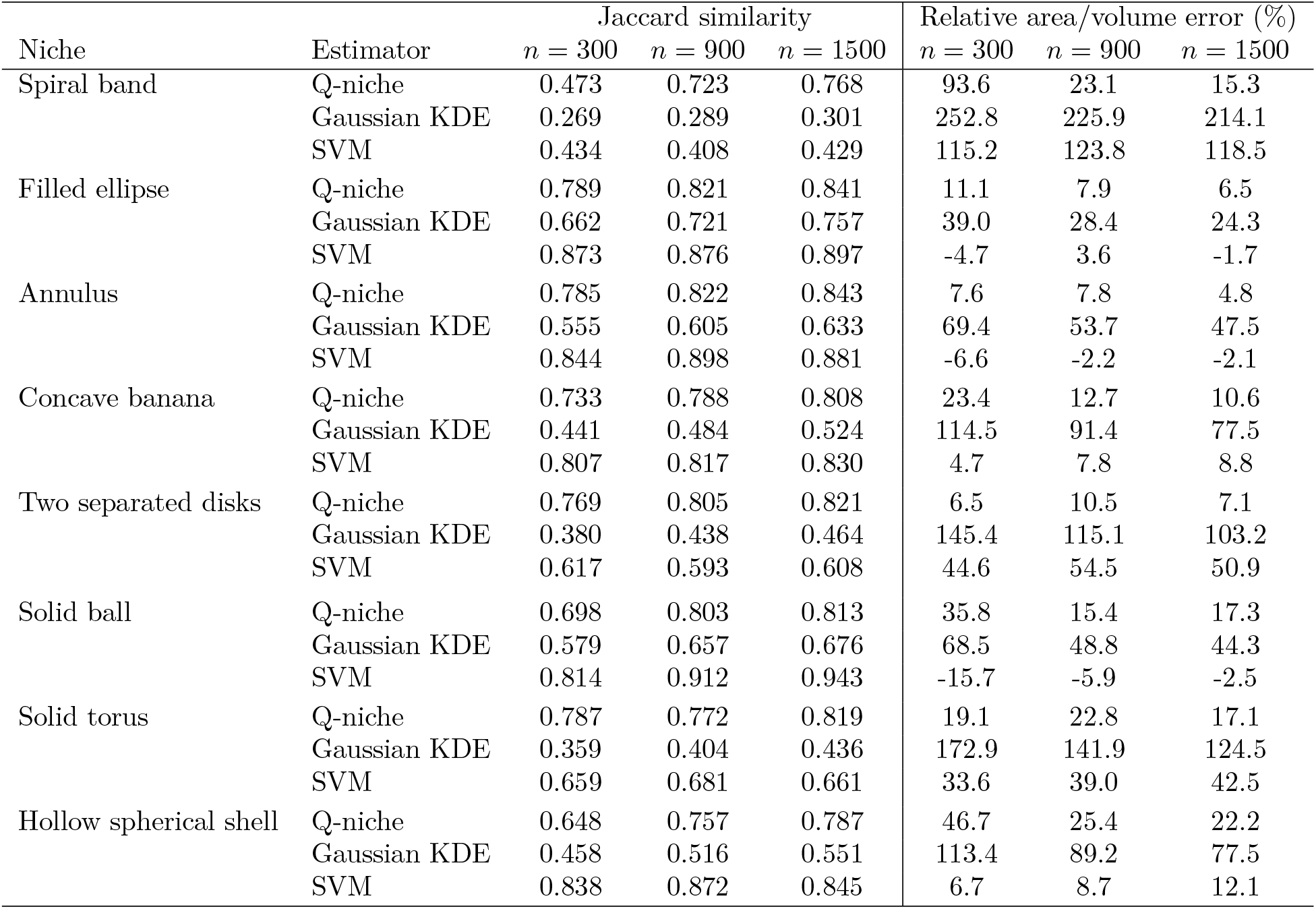
Geometrical recovery of the known synthetic niches across occurrence sample sizes. Jaccard similarity measures overlap between the estimated hypervolume and the known niche, with larger values indicating greater agreement. Relative area or volume error is expressed as a percentage of the analytical niche size; positive values indicate overestimation and negative values indicate underestimation.

| Niche | Estimator | Jaccard similarity |  |  | Relative area/volume error (%) |  |  |
| --- | --- | --- | --- | --- | --- | --- | --- |
| | | $n = 300$ | $n = 900$ | $n = 1500$ | $n = 300$ | $n = 900$ | $n = 1500$ |
| Spiral band | Q-niche | 0.473 | 0.723 | 0.768 | 93.6 | 23.1 | 15.3 |
|  | Gaussian KDE | 0.269 | 0.289 | 0.301 | 252.8 | 225.9 | 214.1 |
|  | SVM | 0.434 | 0.408 | 0.429 | 115.2 | 123.8 | 118.5 |
| Filled ellipse | Q-niche | 0.789 | 0.821 | 0.841 | 11.1 | 7.9 | 6.5 |
|  | Gaussian KDE | 0.662 | 0.721 | 0.757 | 39.0 | 28.4 | 24.3 |
|  | SVM | 0.873 | 0.876 | 0.897 | -4.7 | 3.6 | -1.7 |
| Annulus | Q-niche | 0.785 | 0.822 | 0.843 | 7.6 | 7.8 | 4.8 |
|  | Gaussian KDE | 0.555 | 0.605 | 0.633 | 69.4 | 53.7 | 47.5 |
|  | SVM | 0.844 | 0.898 | 0.881 | -6.6 | -2.2 | -2.1 |
| Concave banana | Q-niche | 0.733 | 0.788 | 0.808 | 23.4 | 12.7 | 10.6 |
|  | Gaussian KDE | 0.441 | 0.484 | 0.524 | 114.5 | 91.4 | 77.5 |
|  | SVM | 0.807 | 0.817 | 0.830 | 4.7 | 7.8 | 8.8 |
| Two separated disks | Q-niche | 0.769 | 0.805 | 0.821 | 6.5 | 10.5 | 7.1 |
|  | Gaussian KDE | 0.380 | 0.438 | 0.464 | 145.4 | 115.1 | 103.2 |
|  | SVM | 0.617 | 0.593 | 0.608 | 44.6 | 54.5 | 50.9 |
| Solid ball | Q-niche | 0.698 | 0.803 | 0.813 | 35.8 | 15.4 | 17.3 |
|  | Gaussian KDE | 0.579 | 0.657 | 0.676 | 68.5 | 48.8 | 44.3 |
|  | SVM | 0.814 | 0.912 | 0.943 | -15.7 | -5.9 | -2.5 |
| Solid torus | Q-niche | 0.787 | 0.772 | 0.819 | 19.1 | 22.8 | 17.1 |
|  | Gaussian KDE | 0.359 | 0.404 | 0.436 | 172.9 | 141.9 | 124.5 |
|  | SVM | 0.659 | 0.681 | 0.661 | 33.6 | 39.0 | 42.5 |
| Hollow spherical shell | Q-niche | 0.648 | 0.757 | 0.787 | 46.7 | 25.4 | 22.2 |
|  | Gaussian KDE | 0.458 | 0.516 | 0.551 | 113.4 | 89.2 | 77.5 |
|  | SVM | 0.838 | 0.872 | 0.845 | 6.7 | 8.7 | 12.1 |

Geometrical agreement generally improved with increasing occurrence sample size, although the magnitude of improvement differed among niches and estimators (Table 2). For Q-niche, increasing sample size substantially improved Jaccard similarity and reduced area overestimation for the spiral band, while relatively low area error and high Jaccard similarity were maintained for the two separated disks across all three sample sizes.

Sensitivity analyses at *n* = 900 showed that geometrical performance varied among estimators and niche shapes, but that the main baseline patterns were generally maintained (Fig. 4). Q-niche Jaccard similarity varied over relatively narrow ranges for most niches. Among the two-dimensional niches, the spiral band showed the greatest variation in both Jaccard similarity and absolute relative area error. Gaussian KDE generally retained lower Jaccard similarity and larger area or volume errors across its tested parameterisations, particularly for the spiral band, separated disks and solid torus. SVM was comparatively stable for the filled ellipse, annulus, solid ball and hollow spherical shell, but showed greater sensitivity for the concave banana, two separated disks and solid torus. Complete baseline and sensitivity results are provided in Table S4.

**Figure 4.**
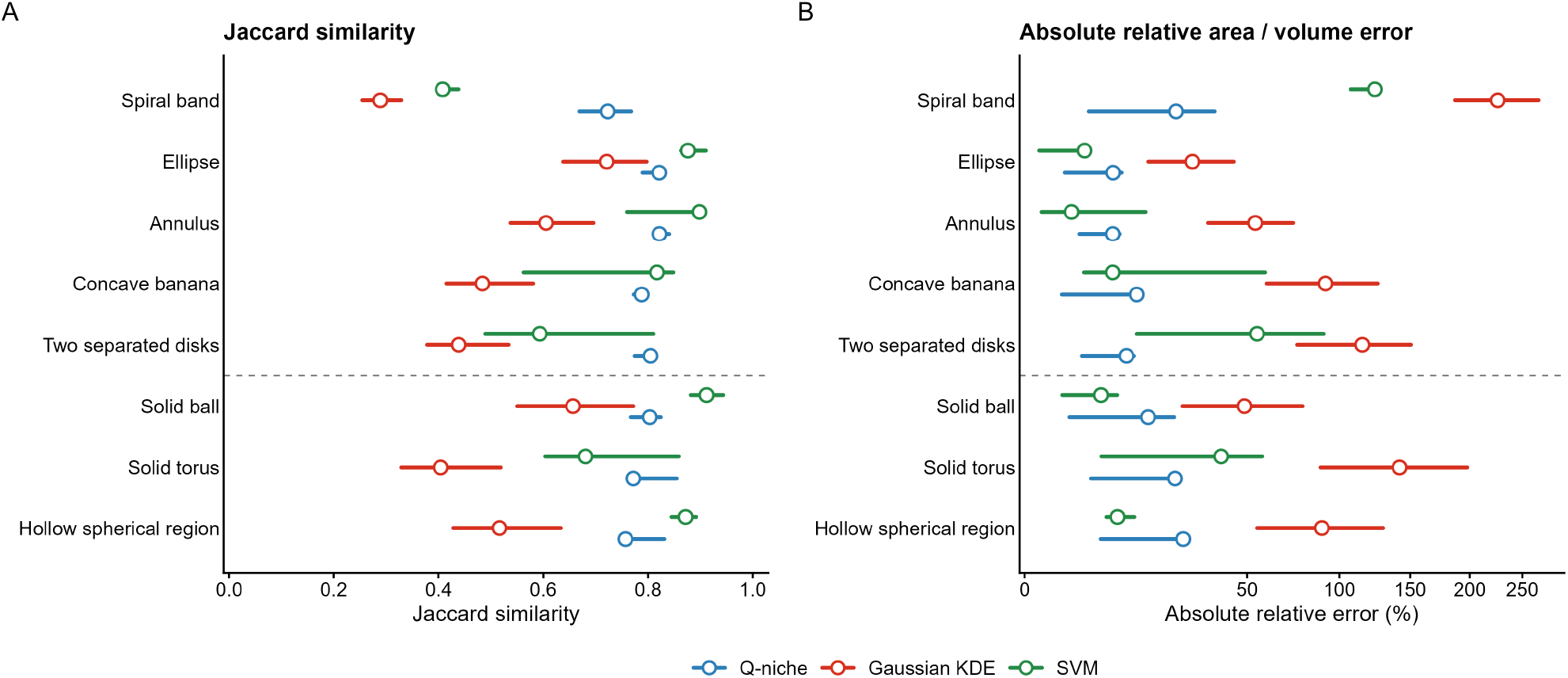
Sensitivity of synthetic niche geometry to estimator parameterisation. Geometrical performance at *n* = 900 is shown for the five two-dimensional and three three-dimensional synthetic niches. (a) Jaccard similarity between each estimated hypervolume and the known niche. (b) Absolute relative area or volume error. Solid coloured lines show the minimum-to-maximum value across the tested sensitivity parameterisations for each estimator, and open circles indicate the corresponding baseline parameterisation. Higher Jaccard similarity and lower absolute relative error indicate closer geometrical agreement with the known niche.

Repeating the two-dimensional baseline analysis at *n* = 900 across 10 independently generated occurrence clouds reproduced the same broad geometrical pattern. Q-niche had the lowest mean absolute relative area error and highest mean Jaccard similarity for the spiral band and two separated disks, whereas SVM performed best for the filled ellipse, annulus and concave banana (Table S3). Across the tested parameter settings, Q-niche showed relatively consistent geometrical performance and achieved the strongest baseline recovery of the spiral band, separated disks and solid torus. SVM more closely reconstructed several other niches, whereas Q-niche tended to produce broader estimates of the known regions; this overestimation contributes to reduced Jaccard similarity. Overall, Q-niche provided a consistent reconstruction of varied niche geometries through a stochastic point-cloud representation suitable for subsequent hypervolume analyses.

### Dimensionality alters estimator behaviour and computational cost

To isolate the effect of dimensionality, we extended the two separated disks into their *d*-dimensional analogue, two separated *d*-dimensional balls, across *d* = 2–7 dimensions at *n* = 900, adjusting ball radius so that total analytical volume remained constant across dimensions. Under the baseline sampling effort of 100 samples per point, Q-niche retained comparatively stable geometrical performance across dimensions, with relative volume error remaining between 6.8% and 30.4% through *d* = 6, while Gaussian KDE increasingly overestimated volume and SVM showed intermediate behaviour. Most notably, Q-niche preserved the expected disconnected *H*_0_ structure in every evaluable analysis from *d* = 2 to *d* = 6, with an exceedance proportion of 1.00 relative to the matched ellipsoid criterion throughout, indicating little evidence that increasing dimensionality itself degraded topological preservation. The main high-dimensional limitation was instead stochastic sampling efficiency: the Q-niche bandwidth increased from *h* = 0.410 at *d* = 2 to *h* = 0.815 at *d* = 7, enlarging the occurrence-centred candidate region while the proportion of sampled points retained by the potential criterion declined sharply. Increasing samples per point mitigated this effect and restored sufficient retained points for persistent-homology analysis at *d* = 6, although even 400 samples per point remained insufficient for the locked 300-point topological subsampling at *d* = 7. Reducing the extent of candidate sampling (corresponding to sd.count in the hypervolume package) as bandwidth increases will also improve candidate acceptance and reduce the sampling effort required, provided that the smaller candidate region still contains the intended potential sublevel set. Q-niche construction time increased with dimensionality and sampling effort. SVM was faster in lower dimensions, but its runtime increased more sharply and exceeded that of Q-niche in six and seven dimensions. Gaussian KDE was faster than both in the highest dimension tested. Full geometrical, topological and runtime results are given in Table S5.

Together, these results demonstrate that Q-niche maintained comparatively stable volume estimates as dimensionality increased and preserved the expected separation between niche components in all assessable cases through six dimensions.

### Virtual species reveal a distinction between environmental topology and geographical overlap

We next moved from analytically controlled niches to two Australian virtual species with known environmental and geographical distributions. Environmental conditions were represented by the first three principal components of 19 bioclimatic variables and elevation. One virtual species had a unimodal environmental response, producing a single connected niche in environmental PC space, whereas the second had a bimodal response whose environmental niche comprised two deliberately disconnected components (see Methods). The resulting suitability surfaces were converted to known geographical presence–absence distributions, from which nested occurrence samples of *n* = 300, 900 and 1500 were drawn and used to construct Q-niche, Gaussian KDE and SVM hypervolumes. We then evaluated recovery separately in environmental and geographical space: environmental overlap measured agreement with the known niche across environmental combinations realised within Australia, whereas geographical overlap measured agreement with the known presence–absence distribution after each estimated hypervolume was projected back across Australia. Environmental Jaccard similarity treated each valid Australian environmental coordinate equally, while geographical Jaccard similarity was weighted by raster-cell area. Higher Jaccard similarity indicates closer agreement with the known niche or distribution. We also assessed inclusion of the deliberately excluded environmental gap and preservation of component separation using the matched ellipsoid criterion. Geographical area error measured overestimation or underestimation of the known distribution, while smaller distances between geographical centroids indicated closer recovery of its position. Complete virtual-species construction and parameterisation are provided in Table S6.

For the unimodal species, SVM produced the strongest overall reconstruction in both environmental and geographical space. It had the highest environmental Jaccard similarity at *n* = 900 and the highest area-weighted geographical Jaccard similarity across all three occurrence sample sizes (Table S7). Q-niche and Gaussian KDE produced similar, broader geographical projections with lower overlap with the known distribution.

The disconnected species revealed a stronger distinction between overall overlap and internal structure (Fig. 5A). At *n* = 900, SVM again produced the highest environmental Jaccard similarity with the known niche, followed by Q-niche and Gaussian KDE (Table S7). The bimodal niche deliberately excluded an intervening region of environmental space between its two suitable components, providing a direct measure of whether estimators interpolated across the gap. At *n* = 900, Q-niche classified 17.3% of environmentally available cells in this region as belonging to the niche, compared with 41.5% for SVM and 77.6% for Gaussian KDE. Persistent homology supported this structural difference: Q-niche had the lowest mean normalised *H*_0_ bottleneck distance from the occurrence cloud. Under the matched ellipsoid criterion, the expected component separation was retained in all ten occurrence repetitions and nine of ten Q-niche repetitions, but in none of the Gaussian KDE or SVM repetitions. By contrast, normalised bottleneck distances for the unimodal species were approximately 0.03 or lower in *H*_0_, *H*_1_ and *H*_2_, indicating substantially smaller topological differences among estimators for the simpler niche.

**Figure 5.**
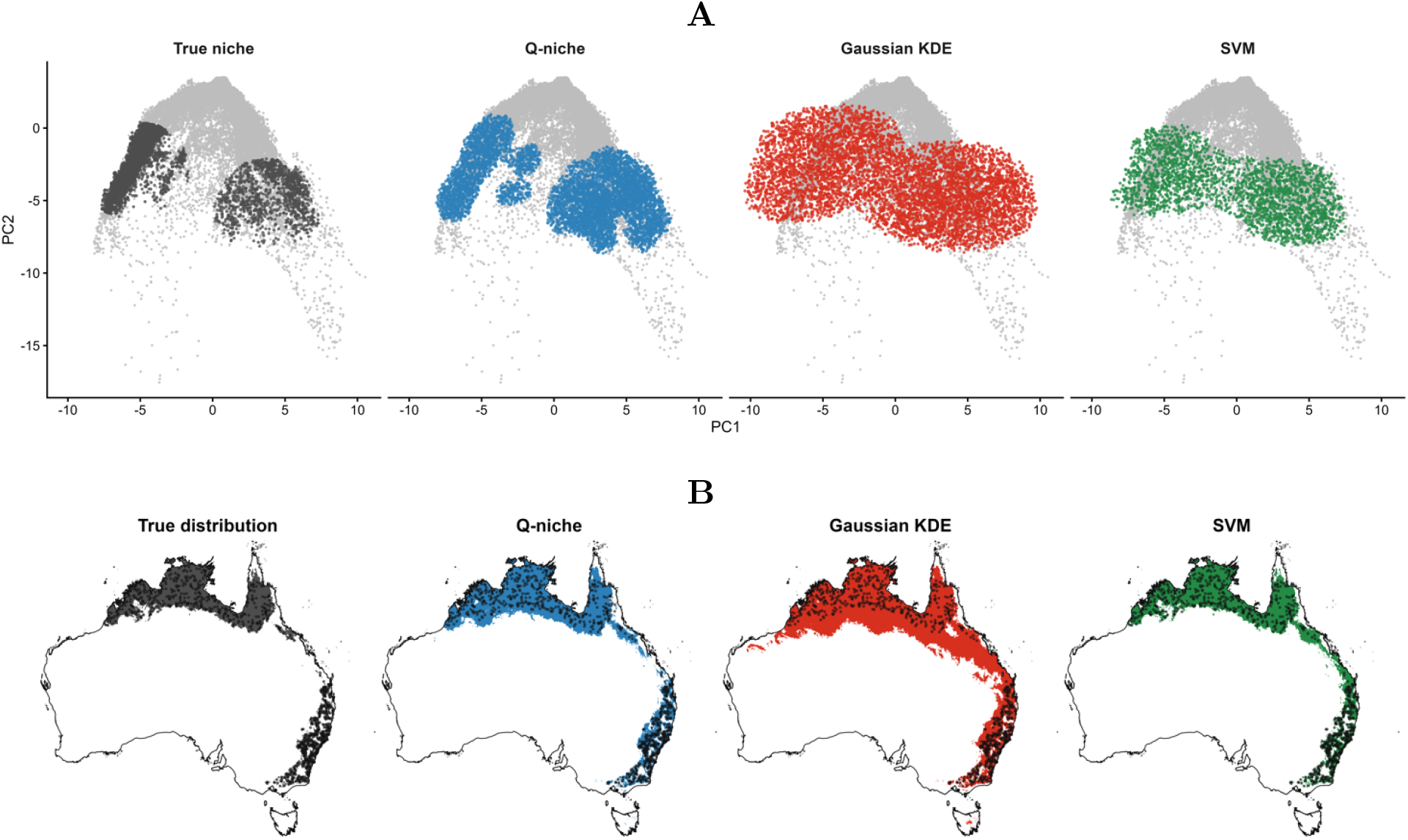
Environmental and geographical reconstruction of a disconnected virtual-species niche. A virtual species comprising two known, disconnected environmental niche components was reconstructed from *n* = 900 occurrence observations using Q-niche, Gaussian kernel density estimation (KDE) and one-class support vector machines (SVM). **(A)** Environmental reconstruction in the PC1–PC2 projection of the three-dimensional environmental niche. Grey points show environmental combinations realised across Australia, and coloured points show the known niche or stochastic points from each fitted hypervolume. **(B)** Corresponding geographical projections across Australia. Coloured raster cells show the known or reconstructed geographical distribution and black points show sampled occurrences.

For the disconnected species, stronger preservation of environmental topology did not translate into greater geographical overlap (Fig. 5B). SVM produced the highest geographical Jaccard similarity at every occurrence sample size and the smallest geographical area error at *n* = 900 (Table S7). All three estimators overestimated geographical extent at *n* = 900, with Q-niche showing less overestimation than Gaussian KDE. Q-niche nevertheless reproduced geographical position most closely: the distance between the area-weighted geographical centroids of the reconstructed and known distributions was 70.4 km for Q-niche, compared with 130.7 km for SVM and 171.9 km for Gaussian KDE. Across the tested Q-niche bandwidth, potential-quantile and sampling-effort settings, geographical Jaccard for the disconnected species ranged from 0.710 to 0.803, while inclusion of the between-mode environmental gap remained between 9.6% and 20.5%.

Together, these results show that Q-niche retained the disconnected environmental structure more consistently than Gaussian KDE or SVM and limited interpolation across the gap between suitable components. This advantage did not translate into the highest geographical overlap, which SVM achieved for both virtual species, although Q-niche most closely recovered the geographical centroid of the disconnected species. Preserving environmental niche topology and recovering geographical extent therefore capture different aspects of estimator performance. Complete environmental, topological, geographical and sensitivity results are provided in Table S7.

### Empirical hypervolume estimates differ in geometry, topology and geographical projection

Finally, we asked whether the estimator-dependent differences observed in the controlled analyses were also apparent in empirical data. We analysed occurrence records for five Australian *Acacia* species, with sample sizes ranging from 414 to 1236 observations, and represented all species within the same three-dimensional PC1–PC3 environmental space. Q-niche, Gaussian KDE and SVM hypervolumes were estimated independently from each occurrence cloud. Geometrical comparisons used hypervolume size, centroid separation and Jaccard overlap, while topological comparisons used normalised bottleneck distances to the occurrence clouds and exceedance proportions under the matched ellipsoid criterion. Higher Jaccard similarity and smaller centroid separations indicated greater agreement among reconstructions, whereas smaller bottleneck distances indicated closer topological agreement with the occurrence clouds. We also compared the total areas of their potential geographical projections across Australia. Because the true environmental niches of these species are unknown, these comparisons describe differences among alternative reconstructions of the same empirical observations rather than estimation accuracy.

The three estimators differed substantially in environmental hypervolume geometry (Fig. 6). All methods ranked *A. saligna* as having the largest environmental hypervolume and *A. georginae* as having the smallest, but the intermediate rankings differed. SVM produced the smallest hypervolume for every species, whereas the relative sizes of Q-niche and Gaussian KDE estimates varied among species. The methods also reconstructed different regions of environmental space: within-species Jaccard similarities between method-specific hypervolumes ranged from 0.327 to 0.582, and their centroids differed in position. Q-niche sensitivity analyses showed that bandwidth and potential-quantile choice could noticeably alter absolute hypervolume size, although species rankings and the pattern of interspecific overlap remained largely stable across the tested settings. Full geometrical comparisons and sensitivity results are provided in Table S8.

**Figure 6.**
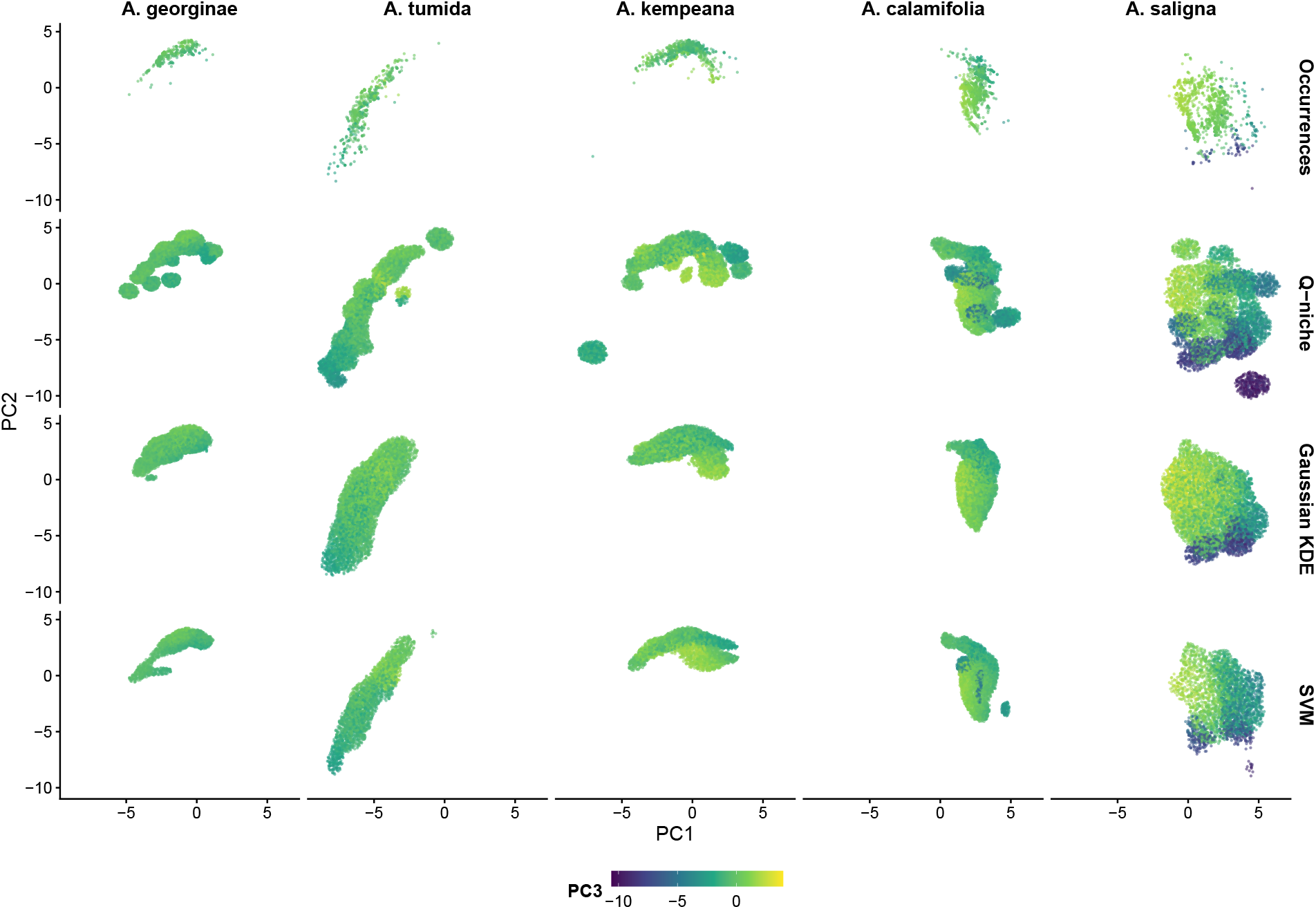
Environmental hypervolume estimates for five empirical *Acacia* occurrence datasets. Occurrence records and environmental hypervolumes estimated using Q-niche, Gaussian kernel density estimation (KDE) and one-class support vector machines (SVM) are shown for *Acacia georginae, A. tumida, A. kempeana, A. calamifolia* and *A. saligna*. All species were represented in the same three-dimensional PC1–PC3 environmental space. Columns correspond to species and rows show the occurrence cloud followed by the Q-niche, Gaussian KDE and SVM estimates. Points are positioned by PC1 and PC2 and coloured by PC3, allowing variation along all three retained environmental dimensions to be visualised.

The matched ellipsoid criterion likewise showed that alternative estimates from the same occurrence cloud could differ in topological structure. For *A. kempeana*, Q-niche consistently showed persistent component separation under this diagnostic, whereas neither Gaussian KDE nor SVM did so. Q-niche also showed consistent component separation for *A. tumida*, while SVM did so less consistently and Gaussian KDE showed no exceedance. The pattern differed for other species: Gaussian KDE showed the greatest *H*_0_ exceedance for *A. georginae*, and SVM did so for *A. calamifolia*. Estimates also differed in *H*_1_ structure, particularly for *A. calamifolia*, while *H*_2_ exceedance was rare across the fitted hypervolumes. Thus, estimator choice affected whether an occurrence cloud was represented by persistent component separation or loop-like structure under the matched ellipsoid diagnostic. Normalised bottleneck distances to the occurrence clouds varied among species, estimators and homology dimensions, with no estimator consistently producing the closest agreement across all comparisons (Table S8).

These differences were also apparent in potential geographical projections (Fig. S1; Table S8). Q-niche produced the largest projected area for four of the five species; Gaussian KDE produced the largest only for *A. georginae*. Within species, the largest and smallest method-specific projections differed by approximately 0.47–1.08 million km^2^. Estimator choice therefore affected both the reconstructed environmental hypervolume and the extent of the associated potential geographical projection. The geographical differences need not be proportional to differences in environmental hypervolume size, because environmental conditions are distributed unevenly across Australia.

Together, these empirical comparisons show that estimator choice substantially changes the inferred size, position and topological structure of a niche, as well as its potential geographical extent, even when the underlying occurrence data are identical. These differences demonstrate that the estimator is a consequential modelling choice when interpreting empirical niche hypervolumes.

## Discussion

Q-niche’s clearest advantage in the controlled benchmarks was its consistent preservation of deliberately imposed non-trivial topology. It retained disconnected components, loops and enclosed voids more consistently than Gaussian KDE or SVM across the tested occurrence sample sizes, while also providing strong geometrical recovery. The same broad two-dimensional pattern was reproduced across independently generated occurrence clouds. Alternative SVM parameterisations improved some reconstructions substantially, showing that estimator performance depends on tuning. However, the topology of an empirical niche is generally unknown, making retrospective selection of a topology-preserving parameterisation difficult. Q-niche therefore appears particularly useful where non-convexity, separation or internal structure may be important but cannot be specified in advance.

The ability to retain such structure matters because observed and realised niches need not be convex or simply connected, and gaps, separated regions and holes may contain information that is lost when an estimator interpolates across them [2, 4, 12, 20]. Blonder proposed that holes within ecological hypervolumes could reflect processes such as exclusion or susceptibility to invasion [12], and subsequent work has developed approaches for detecting topological structure in ecological data [21, 22, 26]. Such features cannot automatically be assigned a biological mechanism: realised niche structure can also reflect environmental availability, accessibility, historical contingency, population structure or incomplete sampling [2, 12, 20, 27]. Nevertheless, investigating whether these structures carry ecological information first requires an estimator that does not systematically remove them. Any continuous hypervolume estimator must infer environmental conditions that were not directly observed in a finite occurrence sample; the relevant question is therefore whether that interpolation removes structure supported by the observations. The contrast between Q-niche and Gaussian KDE is informative in this respect because both use occurrence-centred Gaussian kernels. KDE delineates the niche directly from the smoothed Gaussian density field, whereas Q-niche applies the inverse Schrödinger transformation before defining the retained region. The resulting potential depends on the local curvature of the Gaussian field as well as its magnitude, allowing potential barriers to remain between regions that density smoothing would otherwise connect [28, 29, 36]. The quantum-clustering formulation is useful because it provides this mathematical transformation, not because ecological niches are assumed to be quantum systems.

The virtual-species analyses further showed that better preservation of environmental niche topology did not necessarily produce greater overlap with the known geographical distribution. For the disconnected species, Q-niche retained the separation between the two environmental components much more strongly than the other estimators, whereas SVM produced the highest environmental and area-weighted geographical Jaccard similarity. This distinction is consistent with the Hutchinsonian duality between environmental and geographical space [2, 5, 11]: because environmental conditions are distributed unevenly across geographical space, differences among estimated niches in environmental space need not translate proportionally into geographical overlap. These results suggest a direct ecological application for niche topology. Future virtual-species experiments could manipulate niche topology and environmental availability to test when disconnected components, gaps or holes alter geographical distributions.

The empirical *Acacia* analyses reinforced the conclusion that estimator choice can alter both the geometry and topology inferred from the same occurrence data. The three methods produced substantially different hypervolume sizes, environmental overlap and potential geographical projections, while the matched ellipsoid criterion showed that individual occurrence clouds could be represented as having persistent component separation or loop-like structure under one estimator but not another. Because the true empirical niches are unknown, these differences do not establish which reconstruction is most accurate. They do show why investigators should choose an estimator in light of the structure they consider plausible, the features they wish to retain, and the limitations of their occurrence data. Q-niche also showed appreciable displacement of fitted hypervolume centroids from the corresponding occurrence-cloud centroids, highlighting its sensitivity to environmentally isolated observations. This issue is particularly relevant because observations with large nearest-neighbour distances contribute strongly to the mean squared-distance statistic from which the Q-niche band-width is derived. An isolated observation may therefore influence both the estimated bandwidth and the extent of the reconstructed niche. This is not necessarily undesirable: an environmentally isolated occurrence may represent genuine niche structure that should be retained, but it may instead reflect georeferencing error, environmental-data error or transient occupancy [39].

Across the sensitivity analyses, bandwidth and the potential quantile were the principal determinants of Q-niche extent, whereas stochastic sampling effort had relatively little influence on low-dimensional reconstructions once sufficient points were retained. The dimensionality benchmark provided an important qualification to this pattern. Q-niche continued to preserve the expected disconnected *H*_0_ structure in every evaluable analysis from *d* = 2 through *d* = 6, providing little evidence of deterioration in this diagnostic over the evaluable dimensions. The principal high-dimensional limitation observed was the efficiency of the stochastic representation. The Q-niche bandwidth increased with dimensionality, enlarging the occurrence-centred candidate region while the proportion of candidate points retained by the potential criterion declined sharply. Increasing samples per point restored sufficient retained points for persistent-homology analysis at *d* = 6, but even 400 samples per point were insufficient for the fixed 300-point topological subsample at *d* = 7. Future implementations could improve efficiency through adaptive candidate sampling or other strategies that concentrate sampling within regions likely to satisfy the potential criterion. More generally, the dimensionality used to represent a niche remains an important modelling choice and should be ecologically justified, with its influence on hypervolume geometry, sampling efficiency and computational cost examined explicitly.

No single hypervolume estimator is expected to be optimal for every ecological application. Estimator choice should depend on the structure the investigator wishes to preserve and the assumptions considered reasonable for the dataset. A density-based approach such as Gaussian KDE may be appropriate when smoothing between observations is desired. SVM performed particularly well for several geometrically simpler synthetic niches and for overall overlap with the known virtual-species distributions. Where disconnected components, gaps or holes are themselves of ecological interest, the controlled benchmarks show a clear advantage for Q-niche in preserving that structure. By retaining such features while still reconstructing a continuous niche from finite occurrence data, Q-niche provides a practical route to asking not only how much environmental space a species occupies, but how that space is organised and when its topology matters for ecological and geographical inference.

## Declarations

### Authors’ contributions

JT and JM jointly conceptualised the model and approach presented in the study. JT developed and implemented the Q-niche algorithm, compiled the data, conducted the simulations and analyses, and wrote the initial manuscript draft. Both authors contributed to revising the manuscript and the concepts and approaches. JM was responsible for oversight of the project, project supervision and management.

### Data and code availability

All R code and example data used in this study are publicly available on Zenodo at https://doi.org/10.5281/zenodo.23211950 [40]. The Zenodo archive corresponds to version 1.0.1 of the repository.

## Acknowledgements

JM is supported by the Gordon and Betty Moore Foundation (GBMF13114), the BBSRC (BB/V015249/1), and by the São Paulo Research Foundation (FAPESP) – Grant [Young Investigator: 2025/10520-9]. This research was supported in part by grants from the NSF (DMS-2235451) and Simons Foundation (MPS-NITMB-00005320) to the NSF-Simons National Institute for Theory and Mathematics in Biology (NITMB). The funders had no role in study design, data collection and analysis, decision to publish, or preparation of the manuscript.

## Methods

### Data

#### Synthetic niches

We generated eight analytically defined synthetic niches with known geometry and topology. The two-dimensional benchmark comprised a finite-width spiral, filled ellipse, annulus, concave banana and two separated disks, while the three-dimensional benchmark comprised a solid ball, solid torus and hollow spherical shell. Occurrence points were sampled uniformly from the complete region of each niche. For each geometry, a fixed-seed master cloud of *n* = 1500 observations was generated and nested subsets of *n* = 300, 900 and 1500 were retained; the two-disk subsets contained equal numbers of observations from each component. Exact generating definitions, parameter values, analytical areas or volumes and expected Betti numbers are given in Table S1.

#### Virtual species

We generated two virtual species across Australia using the 19 WorldClim bioclimatic variables and elevation at 2.5-arc-minute resolution [41]. A principal-component analysis was fitted using `virtualspecies::generateSpFromPCA()` to a reproducible sample of 100,000 Australian raster cells, and PC1–PC3 were retained as the common environmental space [42]. The first species had a broad unimodal Gaussian response, which was converted to a deterministic presence–absence distribution using a threshold giving a target geographical prevalence of 0.15.

The second species reused the same PC1–PC3 environmental space but was constructed from two separated Gaussian response modes. The responses were combined by their maximum suitability and their widths reduced until the thresholded environmental supports were non-overlapping, after which the same prevalence criterion was applied. A master set of *n* = 1500 occurrence localities was generated for each species and nested subsets of *n* = 300, 900 and 1500 were retained. Occurrences for the disconnected species were sampled equally from its two environmental components. Exact response parameters used to construct both virtual species are given in Table S6.

#### Empirical *Acacia* data

Occurrence records for *Acacia georginae, A. tumida, A. kempeana, A. calamifolia* and *A. saligna* were obtained from the `acacia_pinus` dataset distributed with the `hypervolume` R package [16, 17]. The package documentation attributes these records to the Botanical Information and Ecology Network (BIEN) database [43]. After environmental-data screening, the analysis retained 414, 578, 806, 1073 and 1236 occurrence records for the five species, respectively. Environmental values were extracted from the same 19 WorldClim bioclimatic variables and elevation used for the virtual-species analyses [41]. Exact coordinate duplicates within species were retained. A single centred and scaled PCA was fitted with `stats::prcomp()` to the pooled occurrence environments of all five species. PC1, PC2 and PC3 explained 40.83%, 34.88% and 9.78% of the environmental variance, respectively, and together accounted for 85.49%; these three axes were retained as the common environmental space for all empirical analyses.

### Hypervolume estimation

Q-niche was implemented using custom R functions following the formulation described above. For each occurrence dataset, Q-niche used a single isotropic Gaussian bandwidth *h*, applied equally across all environmental dimensions. The bandwidth was estimated from local nearest-neighbour distances using a procedure adapted from Nasios and Bors [36]. Briefly, for each occurrence the squared Euclidean distances to its 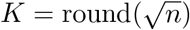 nearest other occurrences were used to calculate a local squared-distance statistic, these values were averaged across observations, and the square root of this average was used as *h*. Full details of the bandwidth calculation are given in Appendix A. Q-niche candidate points were sampled from the union of occurrence-centred sampling regions using the procedure implemented in `hypervolume`, with `sd.count` = 3 and 100 samples per occurrence. Candidate points whose quantum potential was at or below the 0.99 quantile of the occurrence-point potentials were retained, and the resulting point cloud and Monte Carlo volume estimate were stored as a `Hypervolume` object.

All hypervolumes were estimated in R using the `hypervolume` package [16, 17]. Gaussian KDE hypervolumes were fitted using `hypervolume::hypervolume_gaussian()`. Bandwidths were estimated independently for each occurrence dataset using `hypervolume::estimate_bandwidth(method = “silverman”)`, with sd.count = 3 and 100 samples per occurrence; the retained region contained 95% of the estimated probability mass. One-class SVM hypervolumes were fitted using `hypervolume::hypervolume_svm()` with a radial kernel, *ν* = 0.01, *γ* = 0.5, `scale.factor = 1` and 100 samples per point. Fixed random seeds were used throughout, with matched sampling seeds for Q-niche and Gaussian KDE within each dataset.

### Analyses

#### Geometry

Hypervolume geometry was summarised by size, centroid displacement and Jaccard overlap. For the two-dimensional synthetic niches, estimated area and signed percentage error were calculated against the analytical area, and centroid displacement was measured from the known true niche centroid. Jaccard similarity with the known niche was calculated using `hypervolume::hypervolume_set()` with `num.points.max = 10000` and `distance.factor = 1`, using a fixed uniform reference cloud of 100,000 points from each true region. The three-dimensional synthetic analyses similarly compared estimated and analytical volume, calculated centroid displacement from the corresponding fixed true-region reference centroid, and quantified Jaccard and Sørensen overlap with the known niche.

For the virtual species, environmental recovery at *n* = 900 was evaluated over the PC1–PC3 combinations realised by valid Australian raster cells, with each cell contributing equally rather than being weighted by geographical area. We calculated Jaccard similarity with the known binary environmental niche, relative error in the number of occupied environmental cells and Euclidean displacement from the known environmental centroid. For the empirical *Acacia* data, where no true niche was available, we compared estimated hypervolume size, displacement from the corresponding occurrence-cloud centroid, pairwise cross-method Jaccard similarity within each species, and interspecific Jaccard overlap within each estimator. Each fitted hypervolume centroid was calculated as the mean of its stochastic point coordinates in PC1–PC3 space. We also calculated pairwise Euclidean distances between estimator centroids within each species to quantify differences in reconstructed niche position. Empirical geometrical comparisons were therefore treated as differences among alternative reconstructions rather than measures of estimation accuracy.

Geometry sensitivity was assessed at *n* = 900. For the synthetic benchmarks, Q-niche and Gaussian KDE were refitted with bandwidth multipliers of 0.75, 1.0 and 1.25. Q-niche used potential quantiles of 0.95, 0.975 and 0.99, whereas Gaussian KDE used probability quantiles of 0.925, 0.95 and 0.975; stochastic sampling effort was varied over 25, 50, 100 and 150 samples per point for all three estimators. SVM sensitivity was additionally assessed for *ν ∈* {0.005, 0.01, 0.015} and *γ ∈* {0.25, 0.50, 0.75}, varying one parameter at a time. For the virtual species, Q-niche sensitivity used the same bandwidth multipliers, potential quantiles and sampling-effort levels, varying one parameter at a time around the baseline. For the empirical *Acacia* analysis, Q-niche sensitivity used bandwidth multipliers of 0.75, 1.0 and 1.25 and *q ∈* {0.95, 0.975, 0.99}, with sampling effort fixed at 100 samples per point. Complete sensitivity results are given in Tables S4, S7 and S8.

#### Topology

Persistent homology was used to compare topological structure supported by the occurrence clouds with that reconstructed by each hypervolume estimator [22, 44, 45, 46]. Vietoris–Rips persistence diagrams were calculated from Euclidean distances using `TDAstats::calculate_homology()` [47], with `threshold = -1` and no additional coordinate standardisation. The two-dimensional synthetic analyses included *H*_0_ and *H*_1_, whereas the three-dimensional synthetic, virtual-species and *Acacia* analyses included *H*_0_, *H*_1_ and *H*_2_. All comparisons used point clouds of 300 points, with 10 repeated subsamples used to quantify stochastic variation. Only finite persistence features were retained for subsequent comparisons.

Topological agreement was quantified using `TDA::bottleneck()` between each estimator-derived persistence diagram and the corresponding occurrence-cloud diagram. Bottleneck distances were calculated separately for each homology dimension and divided by the diameter, defined as the maximum pairwise Euclidean distance, of the paired occurrence cloud. Values were summarised as mean *±* SD across the 10 repetitions, with smaller normalised distances indicating closer agreement with the topology of the occurrence observations. The occurrence cloud was therefore treated as the topological reference rather than as a complete representation of the true biological niche.

Persistent features were additionally evaluated against matched filled-ellipsoid reference clouds. For each shape or species, sample size and source, 20 reference clouds containing 300 points were generated and transformed to match the realised centroid and covariance matrix of the corresponding full source cloud. For each homology dimension, the matched ellipsoid criterion was defined as the largest finite persistence obtained across the 20 reference clouds; a feature was classified as exceeding the matched ellipsoid criterion only when its persistence exceeded the maximum from all 20 matched ellipsoid clouds. The same threshold was applied to all 10 observed 300-point repetitions; the exceedance proportion was then calculated as the proportion of observed repetitions whose maximum finite persistence exceeded this threshold. In the synthetic analyses, matched ellipsoid clouds were generated for the corresponding occurrence and estimator point clouds, with all two-dimensional synthetic geometries evaluated in *H*_1_ and *H*_0_, and the three-dimensional benchmarks evaluated through *H*_2_. For the virtual species, 20 independent *n* = 300 matched ellipsoid clouds were generated for each species and source from the centroid and covariance of the corresponding source cloud. For the *Acacia* analysis, the same source-level protocol was used, with 20 matched filled ellipsoids generated from the centroid and covariance of each full occurrence or estimator point cloud. Where preservation was classified, a feature was considered retained when both the occurrence and estimator clouds exceeded their respective matched ellipsoid criterion. These comparisons were treated as diagnostics relative to a matched ellipsoid criterion rather than as general tests of statistical or ecological significance.

For the synthetic sensitivity analyses at *n* = 900, bottleneck comparisons were repeated under the parameter perturbations described above using 10 paired 300-point samples. Matched-reference comparisons were also repeated using 20 condition-level matched filled ellipsoids for each occurrence or estimator condition. No additional topological sensitivity analysis was conducted for the virtual-species or empirical Q-niche sensitivity experiments.

#### Geographical projection

Environmental hypervolumes for the virtual species and empirical *Acacia* species were projected geographically across Australia at the native 2.5-arc-minute raster resolution. Environmental values for each valid raster cell were transformed into PC1–PC3 using the same PCA transformation used to construct the corresponding hypervolumes, and each cell was classified as inside or outside the fitted Q-niche, Gaussian KDE or SVM niche. Q-niche inclusion was determined from the fitted potential threshold within its candidate region, Gaussian KDE inclusion from the fitted density threshold, and SVM inclusion from the fitted one-class model. Projections were therefore binary representations of environmental-niche inclusion rather than comparisons of method-specific potential, density or decision scores.

For the virtual species, projected distributions were compared directly with the known binary geographical distributions. Geographical agreement was quantified using cell-area-weighted Jaccard similarity, total projected area and relative area error, and displacement of the area-weighted geographical centroid from the known distribution. For the disconnected virtual species, we quantified inclusion of the deliberately unsuitable environmental gap using a cylindrical corridor between the two niche components, defined in PC coordinates scaled by the Gaussian response standard deviations. Gap inclusion was the percentage of valid Australian raster cells within this corridor classified as belonging to the estimated niche, with each cell weighted equally.

For the empirical *Acacia* species, for which geographical truth was unknown, we compared projected area, area-weighted cross-method Jaccard similarity and the proportion of occurrence localities falling within each projection. Raster-cell areas were calculated using terra::cellSize(). Empirical maps were interpreted as potential geographical projections of the fitted environmental hypervolumes rather than independently validated species distribution models.

For the virtual species, Q-niche geographical sensitivity was evaluated at *n* = 900 by varying bandwidth, potential quantile and sampling effort one parameter at a time around the baseline, using the settings described above (Table S7). For the empirical *Acacia* analysis, geographical projections were also recalculated for the Q-niche sensitivity settings described above.

### Software and reproducibility

Analyses were conducted in R version 4.5.0. Hypervolume estimation used hypervolume version 3.16 [16, 17], and persistent-homology analyses used TDAstats version 0.4.1 and TDA version 1.9.4. Environmental raster processing and geographical analyses used terra version 1.8-60. Q-niche was implemented in custom R code, and fixed random seeds were used for stochastic dataset generation, hypervolume construction, point-cloud resampling and matched-ellipsoid generation. Complete analysis scripts and session information are provided with the accompanying code repository.

## Supplementary materials

**Table S1:** Definitions of controlled synthetic niches. Generating definitions, parameter values, analytical areas or volumes, and expected topological structure for the eight niches used in the two- and three-dimensional synthetic benchmarks.

**Figure S1:** Geographical projections of alternative environmental hypervolume estimates for five *Acacia* species. Environmental hypervolumes estimated using Q-niche, Gaussian kernel density estimation (KDE) and one-class support vector machines (SVM) were projected across Australia using the same environmental variables and principal-component transformation used for hypervolume construction. Columns correspond to *Acacia georginae, A. tumida, A. kempeana, A. calamifolia* and *A. saligna*, and rows correspond to Q-niche, Gaussian KDE and SVM. Coloured raster cells indicate Australian environmental conditions classified as falling within each fitted hypervolume, and black points show the occurrence records used for estimation. These maps are potential geographical projections of the fitted environmental hypervolumes rather than independently validated species distribution models.

**Table S2:** Complete synthetic topology and sensitivity results. Baseline normalised bottleneck distances and matched ellipsoid criterion exceedance proportions, together with topology sensitivity results at *n* = 900, for Q-niche, Gaussian kernel density estimation (KDE) and one-class support vector machines (SVM) across the two- and three-dimensional synthetic niches.

**Table S3:** Robustness of synthetic niche reconstruction to independent occurrence draws at *n* = 900. Results summarise 10 independently generated occurrence clouds for each of the five two-dimensional synthetic niches under the baseline Q-niche, Gaussian KDE and SVM parameterisations.

**Table S4:** Complete synthetic geometry and sensitivity results. Baseline geometrical recovery and one-factor-at-a-time parameter-sensitivity results for Q-niche, Gaussian kernel density estimation (KDE) and one-class support vector machines (SVM) across the two- and three-dimensional synthetic niches. Reported overlap is Jaccard similarity with the fixed true-region reference used for each benchmark.

**Table S5:** Dimensionality benchmark and sampling-effort analysis. Two separated filled *d*-dimensional balls were analysed from *d* = 2 to *d* = 7 at *n* = 900, with total analytical volume and relative component separation held constant across dimensions. Q-niche, Gaussian KDE and SVM were evaluated across 100–400 samples per point.

**Table S6:** Virtual-species construction and parameterisation. Environmental data and PCA construction, virtual-species parameterisation, occurrence-sampling design and baseline estimator settings for the unimodal Gaussian-PCA and disconnected bimodal virtual species.

**Table S7:** Complete virtual-species validation and Q-niche sensitivity results. Environmental and geographical validation, between-mode structural diagnostics for the disconnected species, persistent-homology comparisons, matched ellipsoid criterion results, and the complete one-factor-at-a-time Q-niche sensitivity analysis for the unimodal Gaussian-PCA and disconnected bimodal virtual species.

**Table S8:** Complete empirical *Acacia* hypervolume and Q-niche sensitivity results. Environmental hypervolume size, centroid displacement, cross-method centroid separation and overlap; occurrence-to-estimator normalised bottleneck distances and matched ellipsoid criterion results in *H*_0_, *H*_1_ and *H*_2_; potential geographical projection area and cross-method geographical overlap; and Q-niche sensitivity to bandwidth and potential-quantile choice for five Australian *Acacia* species. Samples per point were fixed at 100 throughout the empirical Q-niche sensitivity analysis.

## Appendix A Quantum-mechanical motivation and mathematical derivation of Q-niche

### The inverse Schrödinger formulation

Q-niche uses the mathematical construction underlying quantum clustering rather than a physical model of ecological systems [28, 29, 36]. Here we provide the quantum-mechanical motivation and complete mathematical derivation of the Q-niche estimator.

In non-relativistic quantum mechanics, the state of a system is represented by a wavefunction *ψ*(**x**, *t*), where **x** denotes position and *t* denotes time. For a normalised wavefunction, its squared magnitude gives the probability density for particle position,

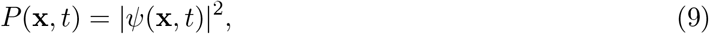

such that *P* (**x**, *t*) d**x** is the probability of observing the particle within an infinitesimal region d**x** around **x** [48].

For a stationary quantum system, separation of the temporal and spatial components gives the time-independent Schrödinger equation

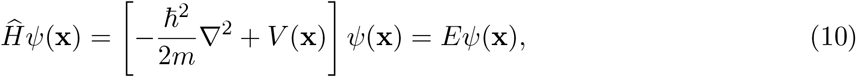

where 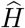 is the Hamiltonian operator, ℏ is the reduced Planck constant, *m* is particle mass, *∇* is the Laplacian, *V* (**x**) is the potential-energy function and *E* is the energy associated with the state *ψ*(**x**) [48]. Conventionally, *V* (**x**) is specified and Equation 10 is solved for the permitted wavefunctions and energies.

Quantum clustering reverses this direction [28, 29, 36]. A smooth positive function constructed from kernels centred on observed data is specified first and used as a stationary wavefunction in the ground-state construction. Rearranging Equation 10 gives

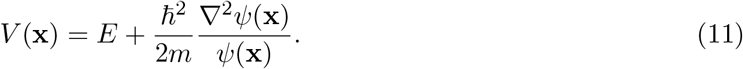

The resulting potential is therefore determined by the local curvature of the specified wavefunction relative to its magnitude.

Q-niche uses this inverse relationship as a dimensionless statistical construction. In the Q-niche formulation, **x** = (*x*_1_, … , *x*_*d*_) denotes an arbitrary point in *d*-dimensional environmental space rather than physical position, and **x**_*i*_ = (*x*_*i*1_, … , *x*_*id*_) denotes the environmental coordinates of occurrence *i*. The constants ℏ and *m* therefore have no ecological interpretation. We instead define bandwidth-standardised environmental coordinates

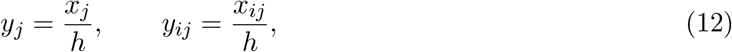

where *h >* 0 is the isotropic Gaussian bandwidth used in every environmental dimension. Thus, *y*_*j*_ is the arbitrary environmental coordinate in dimension *j* expressed in bandwidth units, while *y*_*ij*_ is the corresponding bandwidth-standardised coordinate of occurrence *i*. The inverse Schrödinger construction is then applied directly, with ℏ^2^*/*2*m* set to 1*/*2:

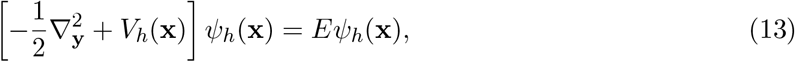

where

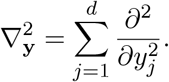

Rearranging gives

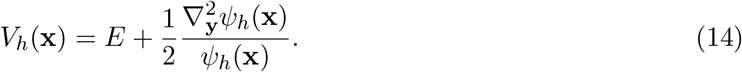

Equation 14 defines the dimensionless inverse Schrödinger relationship used by Q-niche.

### Occurrence-derived Gaussian field

Let

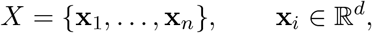

denote the occurrence observations. Q-niche uses a single isotropic Gaussian bandwidth *h >* 0, which is estimated from the local spacing of the occurrence data using a procedure adapted from Nasios and Bors [36].

Nasios and Bors estimate local data spread from the squared Euclidean distances between a focal observation and its *K* nearest neighbours. We retain this local nearest-neighbour construction but adapt the subsequent scale estimator for Q-niche. For a dataset containing *n* occurrences, we set

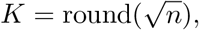

where *K* is the number of neighbouring occurrences used to characterise local spacing around each observation. For each occurrence **x**_*i*_, let **x**_*i*(1)_, … , **x**_*i*(*K*)_ denote its *K* nearest other occurrences, ranked by Euclidean distance. We calculate

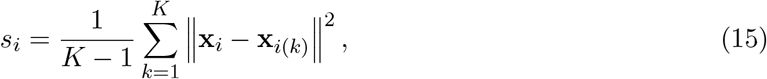

and then average these local squared-distance statistics across all observations,

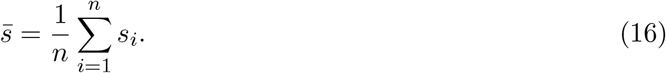

Nasios and Bors model the distribution of these local variance statistics and derive a homoscedastic kernel scale from their mean [36]. Q-niche instead defines the Gaussian bandwidth as

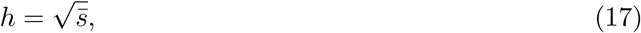

so that the bandwidth has the same units as the environmental coordinates. The same value of *h* is applied in every environmental dimension, giving the isotropic bandwidth vector

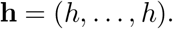

Thus, Q-niche retains the nearest-neighbour measure of local spread introduced by Nasios and Bors while using a square-root transformation and the rule 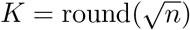 specific to the present estimator.

The squared distance between an arbitrary environmental point **x** and occurrence **x**_*i*_, expressed in bandwidth-standardised coordinates, is

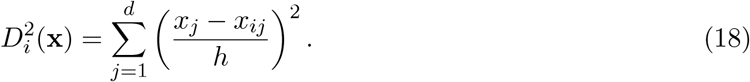

Each occurrence contributes the Gaussian function

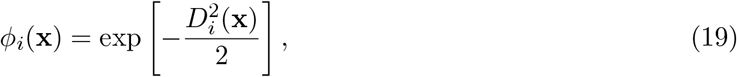

and the complete occurrence-derived field is

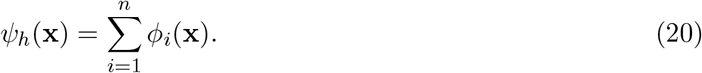

Because every Gaussian contribution is strictly positive, *ψ*_*h*_(**x**) *>* 0 for every finite **x**. The summed field is therefore nodeless, consistent with the positive ground-state construction used in quantum clustering [28, 29, 36]. Q-niche uses this property only as a mathematical device; no physical ground state is attributed to an ecological system.

For an isotropic Gaussian bandwidth *h*, a normalised Gaussian kernel density estimate can be written as

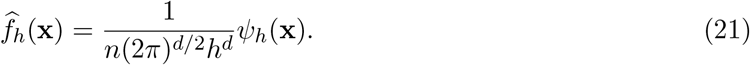

This expresses the algebraic relationship between the occurrence-derived Gaussian field and a Gaussian kernel sum.

### Analytical derivation of the Q-niche potential

To evaluate Equation 14, consider one Gaussian contribution *ϕ*_*i*_. Define

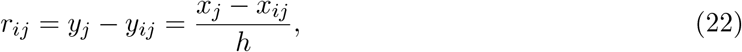

so that

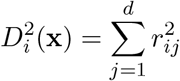

and

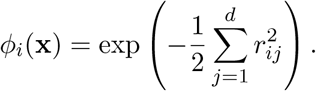

The first derivative of *ϕ*_*i*_ with respect to bandwidth-standardised coordinate *y*_*j*_ is

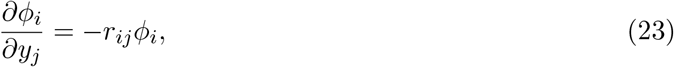

and the corresponding second derivative is

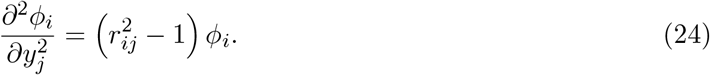

Summing over the *d* environmental dimensions therefore gives

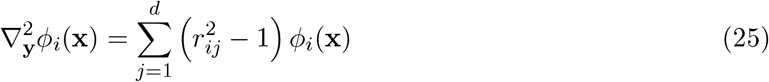

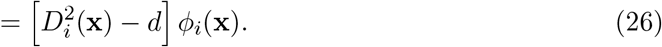

Because differentiation is linear,

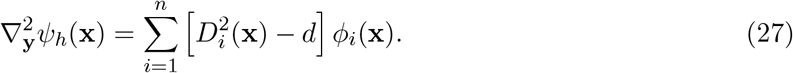

Substituting Equations 20 and 27 into Equation 14 gives

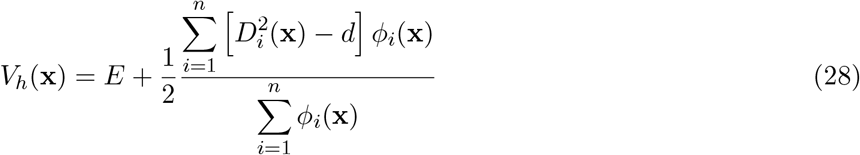

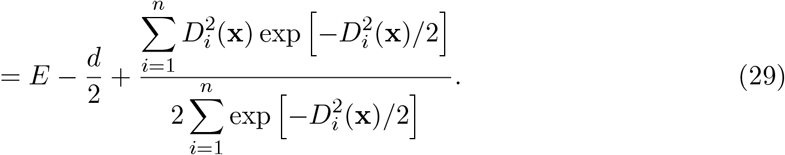

Equation 29 is the analytical Q-niche potential used throughout this study. It can be evaluated directly at arbitrary environmental coordinates and therefore requires neither numerical differentiation nor numerical solution of the Schrödinger equation.

### Interpretation and additive energy constant

The final term in Equation 29 is one half of a Gaussian-weighted mean squared distance from **x** to the occurrence observations in bandwidth-standardised environmental space. Occurrences close to **x** receive greater weight than environmentally distant observations. For a dataset containing a single occurrence,

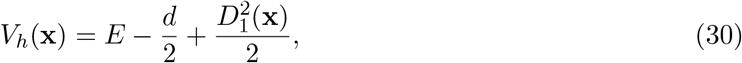

which is a quadratic potential well centred on that occurrence. For multiple occurrences, overlapping Gaussian functions combine to produce a continuous landscape of local wells, ridges and barriers.

The constant *E* produces only a uniform vertical translation of the potential. For any constant *c*,

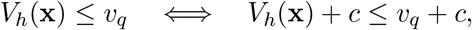

so adding the same constant to the potential and its occurrence-derived threshold leaves the corresponding sublevel set unchanged. We therefore set *E* = 0 in all Q-niche calculations.

### From the potential to the Q-niche hypervolume

The quantum potential is converted into a hypervolume through an occurrence-derived sublevel set. Let

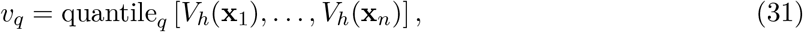

where quantile_*q*_ denotes the empirical *q*th quantile of the occurrence-point potentials, giving the scalar threshold *v*_*q*_. The baseline analyses used *q* = 0.99.

To give the stochastic estimator a finite sampling domain, candidate points are restricted to the union of bandwidth-scaled hyperspheres surrounding the occurrences,

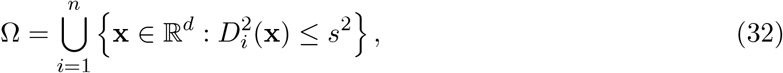

where *s* controls the extent of the candidate region in bandwidth-standardised units. We used *s* = 3 in the baseline analyses, giving hyperspheres of radius 3*h* around each occurrence.

The corresponding Q-niche region is

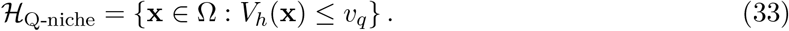

Thus, Q-niche retains environmental conditions that both lie within the finite occurrence-centred candidate region and fall below the occurrence-derived potential threshold.

In the numerical implementation, candidate points **z**_1_, … , **z**_*M*_ were generated approximately uniformly from Ω using the overlap-corrected sampling procedure implemented in the hypervolume package. Define

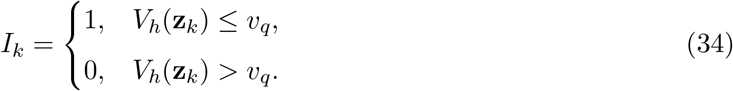

The fraction of the candidate region retained by the Q-niche criterion is estimated as

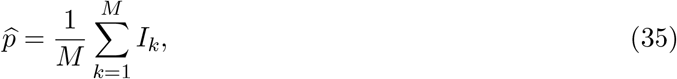

and the Q-niche volume is estimated by

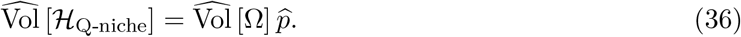

The retained candidate points,

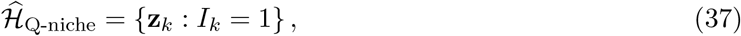

form the stochastic point-cloud representation of Q-niche used in subsequent geometrical, topological and geographical analyses. Increasing the number of candidate points improves the Monte Carlo representation of the same underlying threshold-defined region without changing the analytical potential or its occurrence-derived threshold.

### Interpretation of the Q-niche potential

Q-niche potential values should be interpreted only relatively. Low potential identifies environmental conditions that are more readily retained under the occurrence-derived potential criterion, whereas ridges of higher potential separate lower-potential regions. The potential is not a probability of occurrence, a fitness measure, a physical energy associated with an organism or a direct estimate of physiological suitability.

The topology and geometry of Q-niche therefore arise from the structure of the occurrence-derived Gaussian field, its inverse Schrödinger transformation and the selected sublevel-set threshold. The resulting estimator provides a finite stochastic representation of environmentally supported regions while allowing disconnected components, internal gaps and other non-convex structure to remain represented when supported by the occurrence data.

## References

[1] Jonathan M Chase and Mathew A Leibold. Ecological niches: linking classical and contemporary approaches. University of Chicago Press, 2009.

[2] Jorge Soberón and Miguel Nakamura. “Niches and distributional areas: concepts, methods, and assumptions”. In: Proceedings of the National Academy of Sciences 106.supplement_2 (2009), pp. 19644–19650.

[3] Arnaud Pocheville. “The ecological niche: history and recent controversies”. In: Handbook of evolutionary thinking in the sciences. Springer, 2014, pp. 547–586.

[4] G Evelyn Hutchinson. “Population studies-animal ecology and demography-concluding remarks”. In: Cold Spring Harbor symposia on quantitative biology. Vol. 22. Cold Spring Harbor Lab Press, 1 Bungtown Road, Cold Spring, USA. 1957, pp. 415–427.

[5] Robert K Colwell and Thiago F Rangel. “Hutchinson’s duality: the once and future niche”. In:Proceedings of the National Academy of Sciences 106.supplement_2 (2009), pp. 19651–19658.

[6] George Evelyn Hutchinson et al. An introduction to population ecology. Vol. 260. Yale University Press New Haven, 1978.

[7] Robert H MacArthur. “Population ecology of some warblers of northeastern coniferous forests”. In: Ecology 39.4 (1958), pp. 599–619.

[8] Robert MacArthur and Richard Levins. “The limiting similarity, convergence, and divergence of coexisting species”. In: The american naturalist 101.921 (1967), pp. 377–385.

[9] David Tilman. Resource competition and community structure. 17. Princeton university press, 1982.

[10] Andrew D Letten, Po-Ju Ke, and Tadashi Fukami. “Linking modern coexistence theory and contemporary niche theory”. In: Ecological Monographs 87.2 (2017), pp. 161–177.

[11] Jorge Soberón. “Grinnellian and Eltonian niches and geographic distributions of species”. In:Ecology letters 10.12 (2007), pp. 1115–1123.

[12] Benjamin Blonder. “Do hypervolumes have holes?” In: The American Naturalist 187.4 (2016), E93–E105.

[13] Massimo Pigliucci. “Finding the way in phenotypic space: the origin and maintenance of constraints on organismal form”. In: Annals of Botany 100.3 (2007), pp. 433–438.

[14] William K Cornwell, Dylan W Schwilk, and David D Ackerly. “A trait-based test for habitat filtering: convex hull volume”. In: Ecology 87.6 (2006), pp. 1465–1471.

[15] Christine Lamanna et al. “Functional trait space and the latitudinal diversity gradient”. In:Proceedings of the National Academy of Sciences 111.38 (2014), pp. 13745–13750.

[16] Benjamin Blonder et al. “The n-dimensional hypervolume”. In: Global Ecology and Biogeography23.5 (2014), pp. 595–609.

[17] Benjamin Blonder et al. “New approaches for delineating n-dimensional hypervolumes”. In: Methods in Ecology and Evolution 9.2 (2018), pp. 305–319.

[18] Bassett Maguire Jr. “Niche response structure and the analytical potentials of its relationship to the habitat”. In: The American Naturalist 107.954 (1973), pp. 213–246.

[19] M. P. Austin. “Continuum concept, ordination methods, and niche theory”. In: Annual Review of Ecology and Systematics 16 (1985), pp. 39–61.

[20] M. P. Austin. “A silent clash of paradigms: some inconsistencies in community ecology”. In:Oikos 86.1 (1999), pp. 170–178.

[21] Fernanda S Caron et al. “The evolution of Hutchinsonian climatic niche hypervolumes in gymnosperms”. In: Global Ecology and Biogeography 35.6 (2026), e70257.

[22] Pedro Conceição and Juliano Morimoto. “‘Holey’ niche! finding holes in niche hypervolumes using persistence homology”. In: Journal of Mathematical Biology 84.7 (2022), p. 58.

[23] Andrew L Jackson et al. “Comparing isotopic niche widths among and within communities: SIBER–Stable Isotope Bayesian Ellipses in R”. In: Journal of Animal Ecology 80.3 (2011), pp. 595–602.

[24] Robert R Junker et al. “Dynamic range boxes–a robust nonparametric approach to quantify size and overlap of n-dimensional hypervolumes”. In: Methods in Ecology and Evolution 7.12 (2016), pp. 1503–1513.

[25] Heidi K Swanson et al. “A new probabilistic method for quantifying n-dimensional ecological niches and niche overlap”. In: Ecology 96.2 (2015), pp. 318–324.

[26] Joseph Thomas et al. “Topological data analysis reveals non-convexity in realized niche hypervolumes”. In: bioRxiv (2026). Preprint.

[27] Rodrigo Ramírez-Álvarez, Konrad Górski, and A Townsend Peterson. “A Hierarchical Framework to Assess Ecological Niche Across Levels of Biological Organization”. In: Global Ecology and Biogeography 35.6 (2026), e70258.

[28] David Horn and Assaf Gottlieb. “Algorithm for data clustering in pattern recognition problems based on quantum mechanics”. In: Physical review letters 88.1 (2001), p. 018702.

[29] David Horn and Assaf Gottlieb. “The method of quantum clustering”. In: Advances in neural information processing systems 14 (2001).

[30] David R Nelson and Nadav M Shnerb. “Non-Hermitian localization and population biology”. In: Physical Review E 58.2 (1998), p. 1383.

[31] Karin A Dahmen, David R Nelson, and Nadav M Shnerb. “Life and death near a windy oasis”. In: Journal of mathematical biology 41.1 (2000), pp. 1–23.

[32] J Christopher D Terry, Jacob D O’Sullivan, and Axel G Rossberg. “Schrödinger’s Range-Shifting Cat: How Skewed Temperature Dependence Impacts Persistence with Climate Change”. In: The American Naturalist 203.2 (2024), pp. 161–173.

[33] Raimundo Real, A Márcia Barbosa, and Joseph W Bull. “Species distributions, quantum theory, and the enhancement of biodiversity measures”. In: Systematic Biology 66.3 (2017), pp. 453–462.

[34] Ricardo A Rodríguez et al. “Empirical clues about the fulfillment of quantum principles in ecology: potential meaning and theoretical challenges”. In: Ecological Modelling 301 (2015), pp. 90–97.

[35] Maxime Clenet, Maxime Dion, and F Guillaume Blanchet. “Addressing ecological challenges from a quantum computing perspective”. In: Methods in Ecology and Evolution 17.3 (2026), pp. 632–649.

[36] Nikolaos Nasios and Adrian G Bors. “Kernel-based classification using quantum mechanics”. In: Pattern Recognition 40.3 (2007), pp. 875–889.

[37] Robert Ghrist. “Barcodes: the persistent topology of data”. In: Bulletin of the American Mathematical Society 45.1 (2008), pp. 61–75.

[38] Robert W Ghrist. Elementary applied topology. Vol. 1. Createspace Seattle, 2014.

[39] Xiao Feng et al. “A checklist for maximizing reproducibility of ecological niche models”. In:Nature Ecology & Evolution 3.10 (2019), pp. 1382–1395.

[40] Joseph Thomas. j-thomas4590/Q-niche: Q-niche v1.0.1. Version v1.0.1. Oct. 2026. doi:10.5281/zenodo.23211950. url: https://doi.org/10.5281/zenodo.23211950.

[41] Stephen E Fick and Robert J Hijmans. “WorldClim 2: new 1-km spatial resolution climate surfaces for global land areas”. In: International journal of climatology 37.12 (2017), pp. 4302–4315.

[42] Boris Leroy et al. “virtualspecies, an R package to generate virtual species distributions”. In:Ecography 39.6 (2016), pp. 599–607.

[43] Brian S Maitner et al. “The bien r package: A tool to access the Botanical Information and Ecology Network (BIEN) database”. In: Methods in Ecology and Evolution 9.2 (2018), pp. 373–379.

[44] Herbert Edelsbrunner, John Harer, et al. “Persistent homology-a survey”. In: Contemporary mathematics 453.26 (2008), pp. 257–282.

[45] Steve Y Oudot. Persistence theory: from quiver representations to data analysis. Vol. 209. American Mathematical Society Providence, RI, 2015.

[46] Nina Otter et al. “A roadmap for the computation of persistent homology”. In: EPJ data science 6.1 (2017), p. 17.

[47] Raoul R Wadhwa et al. “TDAstats: R pipeline for computing persistent homology in topological data analysis”. In: Journal of open source software 3.28 (2018), p. 860.

[48] David J Griffiths. Introduction to quantum mechanics. Pearson education, 1995.

